# Prediction of TdP Arrhythmia Risk Through Molecular Simulations of Conformation-specific Drug Interactions with the hERG K^+^, Na_V_1.5, and Ca_V_1.2 Channels

**DOI:** 10.1101/2025.09.25.678690

**Authors:** Kyle C. Rouen, Kush Narang, Yanxiao Han, David Wang, Ensley Jang, Sophia Brunkow, Vladimir Yarov-Yarovoy, Alexander D. MacKerell, Igor Vorobyov

## Abstract

Unintended block of cardiac ion channels, particularly hERG (K_V_11.1), remains a key concern in drug development as disruption of ion channel function can lead to deadly arrhythmia. To assess proarrhythmic risk, we investigated how drugs interact with hERG in its open and inactivated states and whether drug interactions with other cardiac channels like Na_V_1.5 and Ca_V_1.2 mitigate that risk. Using cryo-EM structures, we modeled open and inactivated conformations of these channels with Rosetta and AlphaFold. We then applied Site Identification by Ligand Competitive Saturation (SILCS), a physics-based pre-computed ensemble docking method, to predict drug binding affinities. SILCS leverages molecular simulation-generated free energy maps for high-throughput docking against hydrated lipid bilayer-embedded ion channel models. Bayesian machine learning was used to refine SILCS scoring using experimental IC_50_ values from 69 known hERG blockers outperforming Schrödinger Glide, AutoDock Vina, and OpenEye FRED drug docking predictions. Computed drug binding affinities for hERG and Ca_V_1.2 channels were used to train machine learning models that successfully classified around 300 drugs from the CredibleMeds database. Cationic nitrogen SILCS fragment free energy scores were found to be top physical properties that are predictive of drug-induced Torsades de Pointes (TdP) arrhythmia risk. This approach, which relies on the predicted binding free energies and predicted physical properties of drugs rather than the chemical structure of the drugs themselves as features could be extended to facilitate the design of new drugs where rapid assessment of arrhythmia risk can be performed prior to experimental testing.

## 1. INTRODUCTION

Drug interactions with cardiac ion channels can promote deadly arrhythmia. The hERG K^+^ channel is a critical anti-target in small molecule drug design. By facilitating the rapid component of the delayed rectifier K^+^ current in cardiac myocytes known as *I_kr_*, the hERG channel plays a major role in ending the cardiac actional potential and allowing cardiac myocytes to return to the resting membrane potential^2^. Drug binding to the pore of the hERG channel reduces the repolarizing K^+^ current and can thereby prolong the action potential duration, termed QT prolongation. Drugs such as astemizole, cisapride, and terfenadine have been withdrawn from markets in several countries as a result of QT prolongation resulting from hERG inhibition^3^. In many instances, drug-induced QT prolongation leads to the development of life-threatening Torsades de Pointes (TdP) arrhythmia, a type of ventricular tachycardia. Despite this, many potent hERG blockers present relatively little TdP risk, e.g., an antihypertensive drug verapamil, which blocks hERG in the low nanomolar range.^4^

*In vitro* screening for TdP risk is costly and time-consuming. Furthermore, typical screening protocols fail to capture state-dependent binding affinities^5^. Alternatively, the availability of the cryo-EM structure of the open state of the hERG channel^6^ has spurred a wave of attempts at structure-based drug classification^7^. However, many diverse small molecules bind to the hERG channel^8^, thereby making it challenging to develop computational models for TdP risk prediction. To date, there has been no clear consensus pharmacophore model for a hERG blocker despite multiple efforts to analyze the commonalities of hERG blockers in conjunction with the available structure of the hERG channel ^9, 10^. Our laboratory previously developed a proof-of-concept virtual pipeline that was applied for prediction of pure hERG blocking drugs without off-target effects^11^. While partially successful, this study was limited by considering only the open state of the channel and, therefore, may be limited by state-specific drug binding to the hERG channel that may contribute to proarrhythmic tendencies.

We hypothesized that specific block of the inactivated conformation of the hERG channel leads to a greater incidence of arrhythmia than drugs that bind to the open state based on previous studies^11, 12^. These state-specific interactions are likely to be the critical determinants of drug associated safety or proarrhythmia^13–15^. Previous publications from our laboratory^11, 16^ supported this conclusion: the drug moxifloxacin does not block the inactivated state of hERG ^17, 18^ and is predicted to be safer than dofetilide, confirmed by clinical data^19, 20^. Here, we have generalized this approach beyond these two example drugs.

hERG block alone is not sufficient to predict proarrhythmic risk. Many drugs block the hERG channel but do not significantly promote adverse cardiac events. As mentioned above, verapamil, for instance, remains a common antihypertensive despite having nanomolar affinity for the hERG channel ^4^. Consideration of simultaneous interactions with the Na_V_1.5 and Ca_V_1.2 channels will likely lead to better predictions of the proarrhythmic risk of drugs^1^. Furthermore, Na_V_1.5 channels conduct both early and late sodium currents; drugs that selectively reduce late sodium currents may better protect against the effects of hERG block as compared to other sodium channel blockers^21^. Consequently, we must consider multiple conformational states of each ion channel target to adequately assess the effects of a drug on the cardiac action potential.

Recently, Escobar et al. used different voltage protocols to verify a Markovian model of state-dependent interactions with the hERG channel^22^. This approach of modeling interactions with open, inactivated states in addition to drug trapping in the closed state has proven to be better than simple scaling of *I*_Kr_ current at reproducing the effects of hERG-blocking drugs. Other *in silico* studies use experimental ion channel data to predict features of the cardiac action potential.^23, 24^ Experimental biomarkers have further been used to train *in silico* models.^25, 26^ We show here that this is not necessary. One can achieve comparable performance considering only the chemical structure of the drug and the structures of three protein targets.

Here we use available experimental cryo-EM structures of hERG, Na_V_1.5 and Ca_V_1.2 channels ^6, 27^ with computational modeling methods SILCS (Site Identification by Ligand Competitive Saturation) ^28^ and Rosetta ^29^ to assess the binding of 53 hERG-blocking drugs from two recent studies. ^1, 5^ This is then extended to a larger testing set of 300 drugs from the CredibleMeds database (https://www.crediblemeds.org/). This pipeline, as presented in Figure 1, takes predicted binding free energies and physical-chemical properties about drug molecules to make accurate predictions about clinical arrhythmia risk. While previous work has relied on experimental inputs, our classifier achieves similar performance with only computational predicted binding free energies. In this way, we successfully demonstrate a low-cost and efficient method to screen for cardiac-safe drugs.

**Figure 1:**
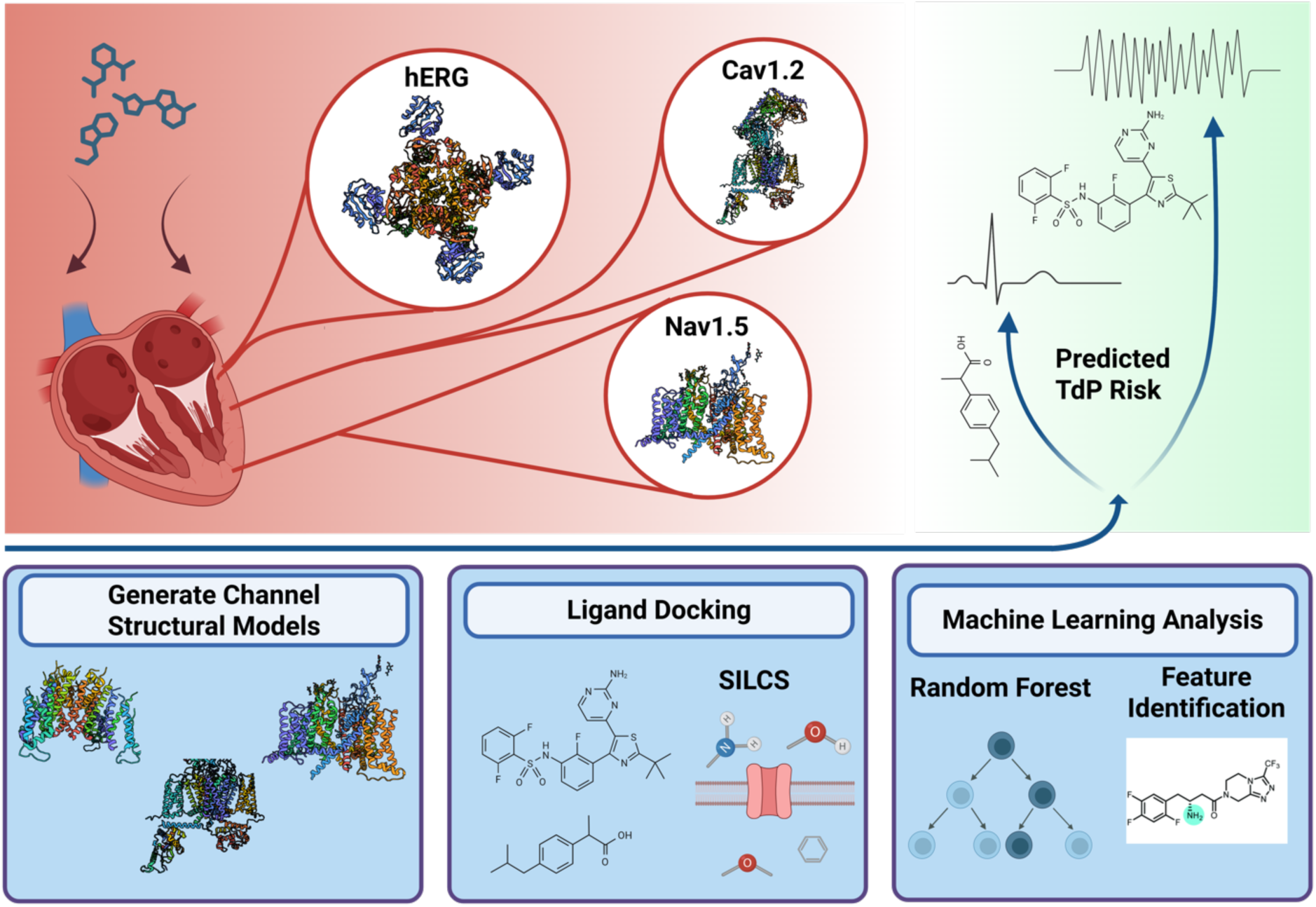
Overview of Cardiotoxicity Prediction Pipeline.

## 2. METHODS

### 2.1. Ion Channel Structures

hERG channel models: We generated structural models of the hERG channel based on available experimental structures deposited in the protein data bank (PDB)^30^. The cryogenic electron microscopy (cryo-EM) structures of the hERG channel in the potentially open conformation from Wang and MacKinnon (PDB IDs 5VA2 and 5VA1)^6^ have been used for most modeling studies. In the same study, Wang and MacKinnon published the cryo-EM structure of the non-inactivating mutant of the hERG channel, S631A (PDB ID 5VA3). A structure of the hERG channel solved under low 3 mM [K^+^] (PDB 9CHQ) represents a putatively more inactivated state of the hERG channel with V625 flipped outward, disrupting the selectivity filter (SF) S2 K^+^ ion binding site.^31^ Here, we rebuilt the missing extracellular loops of 5VA2, 5VA3, and 9CHQ using Rosetta loop modeling to have a more complete structure for subsequent ligand docking.^32^ Visual comparison of the hERG structures and details of the selectivity filter of each channel are shown in Figure S1.

Additionally, we utilized Rosetta structural modeling^32^ to introduce the S641A mutation that promotes channel inactivation^33^. To relax the mutated protein, we simulated the S641A model for 90 nanoseconds using all-atom molecular dynamics (MD) simulations using Amber PMEMD program^34^. The latest-generation CHARMM36m^35^ protein force field based on an earlier foundational CHARMM22 protein force field model developed by A. MacKerell, M. Karplus and co-workers,^36^ the CHARMM36 (C36) lipid force field,^37^ and the CHARMM TIP3P water model^38^ were used in these and other MD simulations in this study. The MD equilibration steps were conducted at 310 K and 1 bar pressure with a 2 fs time step under the periodic boundary condition using the Langevin dynamics thermostat with randomized initial velocities and the Monte Carlo barostat to allow for system volume relaxation. Lennard-Jones interactions were smoothly switched off at 10– 12 Å by a force-switching function with the electrostatic interactions treated using particle Mesh Ewald^39^ with a 12 Å real space cutoff. A lipid membrane composed of 1-Palmitoyl-2-oleoylphosphatidylcholine (POPC) and cholesterol (in a 9:1 ratio) was used and solvated by aqueous 0.15 M KCl. Restraints were initially applied to the alpha carbons of the protein and successively removed as previously described^11^ with all restraints being removed after 40 ns.

Sodium channel models: We utilized two sodium channel structure for ligand docking. First, we used the quinidine-bound and inactivated structure of the human Na_V_1.5 (PDB ID 6LQA)^40^. The homology model from Nguyen et al. based on PDB ID 5XSY (a cryo-EM structure of the electric eel Na_V_1.4 in an inactivated conformation) was also used.^41^

Calcium channel models: We made use of three models of the Ca_V_1.2 channel. First, we used the cryo-EM structure of hCa_V_1.2 with amiodarone/sofosbuvir-bound in an inactivated state (PDB ID 8FHS)^42^. Second, we used the inactivated structure with gabapentin bound to the auxiliary α_2_**δ** subunit (PDB ID 8FD7)^43^. Third, we used an open-state Ca_V_1.2 model built using Rosetta Comparative modeling^44^ based on an open-inactivated state structure of Na_V_1.4 with a slightly widened pore (PDB ID 6AGF)^45^. (See Figure S6 for comparison of pore diameter of Ca_V_1.2 models.)

### 2.2. Ligand Docking

To analyze drug interactions with the hERG channel, we utilized the SILCS methodology. SILCS is a pre-compute ensemble docking software based on grand canonical Monte Carlo (GCMC) and MD simulations^46^. SILCS involves GCMC/MD simulations of the proteins in a collection of 8 solutes (methylammonium, propane, benzene, dimethyl ether, formamide, acetate, acetaldehyde, imidazole) representing most common functional groups of small-molecule ligands at ∼0.25 M in explicit water from which functional group free energy patterns, termed grid free energy (GFE) FragMaps, around the protein are calculated. Since this method includes an explicit lipid membrane, water molecules, and flexibility of the protein, SILCS is more rigorous than other docking methods that rely on implicit membrane and solvent representations and static protein structures^47^. SILCS Monte Carlo (SILCS-MC) docking is used to perform drug docking in the field of the GFE FragMaps and compute ligand binding free energies (LGFE) that can be converted to dissociation constants, *K*_D_. A Bayesian machine learning approach may then be used to improve correlation with experimental IC_50_ values^48^ as was done previously for an open-state hERG channel model. ^49^

SILCS FragMap generation was performed using SILCS 2020.1, 2023.1, and 2024.1 software suite (SilcsBio LLC). Each of the ion channel structural models described above were embedded in lipid membranes composed of 9:1 ratio of POPC lipids to cholesterol built using CHARMM-GUI web toolkit^50^, which runs CHARMM molecular simulation software^51–53^ in the background. Then these molecular systems were solvated by aqueous solutions containing 8 solutes at ∼0.25 M concentration using SILCS scripts. Then 10 independent GCMC/MD simulations of these systems commenced using different initial solute positions and atomic velocities. MD simulations lasted 100 ns each and were intervened every 1 ns by 200,000 GCMC steps. The alpha carbons of protein residues were restrained (0.12 kcal/mol/Å^2^) throughout the GCMC/MD simulations to maintain the overall protein conformation. Models were generally built without Na^+^, K^+^ or Ca^2+^ ions or other small molecules. One set of simulations was run with K^+^ ions restrained in the selectivity filter of the hERG channel at the S0, S2, and S4 sites, consistent with the K^+^ ions observed in the cryo-EM structure 7CN0^27^. SILCS GCMC/MD simulations used the same MD simulation parameters as described above as well as CHARMM36m protein^35^, the C36 lipid, the CHARMM General Force Field^54^ and the CHARMM TIP3P water model^38^ with oscillating chemical GCMC code^55^, the GROMACS simulations package^56^ and the SILCS-MC docking utilities^28^ (SilcsBio LLC). SILCS-MC docking was performed with SILCS version 2024.1. Bayesian machine learning (BML) parameter optimization was used to improve the ligand docking scores compared to experimental IC_50_ values, as described in Goel et al ^46^. A search radius of 10 Å was used for exhaustive SILCS-MC docking. SILCS binding LGFE scores were used as features as well as atom-type specific GFE scores for methylammonium nitrogen (MAMN), benzene carbon (BENC), propane carbon (PRPC), methanol oxygen (MEOO), generic hydrogen bond acceptor (GENA), and generic hydrogen bond donors (GEND).

For all docking to the hERG channel, the search area was centered below the selectivity filter near Tyr652, consistent with the experimentally-identified drug binding site.^57, 58^ Docking to Ca_V_1.2 was performed in a search radius defined based on the center of mass of the amiodarone coordinates from PDB 8FHS. Docking to Na_V_1.5 was performed in a search radius defined based on the center of mass coordinates for quinidine from PDB 6LQA.

Docking was also performed with FRED, an OpenEye software.^59^ Ligands were prepared with OpenEye OMEGA^60^, setting the maximum number of conformers per ligand to 200. During the receptor generation step for the hERG channel models, an outer contour was set to encompass the known binding site below the selectivity filter, encompassing the S624, Y652, and F656 residues.

Docking was further performed with Glide software from Schrödinger^61^ using Standard Precision (SP). The ligands were prepared using LigPrep. For the hERG channel models, Receptor Grid generation was again centered on Y652.

Finally, docking was further performed with AutoDock Vina (version 1.2)^62^. The docking grid was centered at Y652 with a 10 Å search box. Optimized empirical parameters from Pham et al. were initially used to perform docking.^63^ Subsequently, these same empirical parameters were varied to increase the Pearson Correlation of the training set data (See Figure S10).

Ligand structure files were downloaded from PubChem database^64^ and prepared with Open Babel^65^ in the protonation state most abundant at physiological pH. Ligands were collected from Kramer et al.^1^ and Crumb et al studies^5^. The IC_50_ values from these two studies were used as the experimental reference for each drug. Additional drugs were taken from the CredibleMeds database, using the drugs with “Known Risk” as TdP+ and “Therapeutic Options” drugs as TdP-.

### 2.3. Molecular Descriptors

Molecular descriptors for each drug molecule were generated using RDKit^66^. These features include the physical properties such as the water-octanol partition coefficient (LogP), the topological polar surface area (TPSA), and molecular weight. Additionally, we included descriptors relating to the complexity of the molecular structure such as the Balaban J Index (BalabanJ)^67^, the Bertz Complexity Index (BertzCT)^68^, and the Kappa Shape Indices (K1, K2, K3)^69^

### 2.4. TdP Risk Classifiers

Logistic regression analysis was performed using R version 4.3.3^70^ and Python 3.9. A generalized linear model was built (using the *glm()* function) to determine regression coefficients. The *sklearn* package was used to build and assess the logistic regression model. TdP risk classification was taken from Kramer et al. for most drugs.^1^ Additional classifications were based on entries from CredibleMeds. Models were assessed based on the Matthew’s Correlation Coefficient (MCC) and area under the receiver operating characteristic curve (ROC AUC). Optimal thresholds of the predicted probability from each regression model are calculated such that the ROC AUC is maximal. The logistic regression models have the general form that for a given drug, *X*, the log-likelihood of TdP arrhythmia is given by:

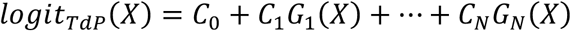

Here, *C*_0_ is a constant intercept term. *G*_N_(*X*) is the Δ*G*_bind_ for drug *X* to the protein target *N* and *C*_N_ is the logistic regression coefficient associated with the protein target *N*. If *C*_N_ < 0, strong drug binding to protein target *N* is associated with greater TdP risk in the model. Conversely, if *C*_N’_ > 0 then that indicates drug binding to protein *N*′ is protective against TdP arrhythmia. Coefficients for the various models are listed in Figure S11.

The random forest classifier was accessed from *sklearn* package.^71^ The XGBoost Classifier was accessed from the *xgboost* package.^72^ The neural network model was generated with PyTorch.^73^ The hyperparameters of each model were tuned using the *optuna* package.^74^ Feature importance to model classification was assessed using SHapley Additive exPlanation (SHAP) value analysis.^75^ All classifier code is available on GitHub: https://github.com/kylerouen/tdp_classifiers.

## 3. RESULTS

### 3.1. Comparison of docking methods

First, we sought to evaluate the use of computational docking to predict the affinity of small-molecule drugs to the cardiac ion channels hERG and Ca_V_1.2. Four different docking methods were tested including SILCS, GLIDE, FRED, and AutoDock Vina. Each docking method gives a docking score in its own internal units, each being roughly proportional to the free energy of binding (Δ*G*_)*+$_). The docking scores were compared to experimental IC_50_ values for each of the 53 drugs from Kramer et al. since all of these drugs were assessed via electrophysiology in the same lab under consistent experimental conditions^1^. The Pearson coefficient was consistently better for docking to the hERG channel (Pearson r ∼ 0.4) as compared to Ca_V_1.2 (Pearson r ∼ 0.2), potentially because the hERG channel binding site is narrower and therefore easier for the docking programs to sample (hERG 5VA2 has a cavity volume of 3730 Å^3^ as compared to Ca_V_1.2 8FD7 with 7391 Å^3^ as measured by OpenEye make receptor software). Notably, OpenEye FRED performed very well with hERG affinity but yielded no significant correlation when docking to Ca_V_1.2 (Figure 2). While all Pearson and Spearman r values for all 4 docking methods were less the 0.5 with all 4 docking methods, SILCS LGFE scores produced reasonably high correlations for both hERG and Cav1.2. However, nifedipine and nitrendipine, two very potent calcium channel blockers (CCBs) with nanomolar affinities, scored poorly with SILCS (Figure S5) as well as for all of the docking programs having the lowest affinity docking scores among all the drugs tested (not shown and Figure S15). In an effort to improve docking results, docking was confined to fenestration sites, consistent with cryo-EM structure of the nifedipine-bound rabbit Ca_V_1.1 channel. The RMSD of the docking pose was significantly improved, but this resulted in no improvement in the correlation between docking scores from FRED and experimental affinity values (Figure S15). The failure of the docking programs to identify potent CCBs represents a significant limitation when assessing the cardiac safety of drug molecules as strong calcium channel block can significantly counteract the effect of hERG channel block.

**Figure 2:**
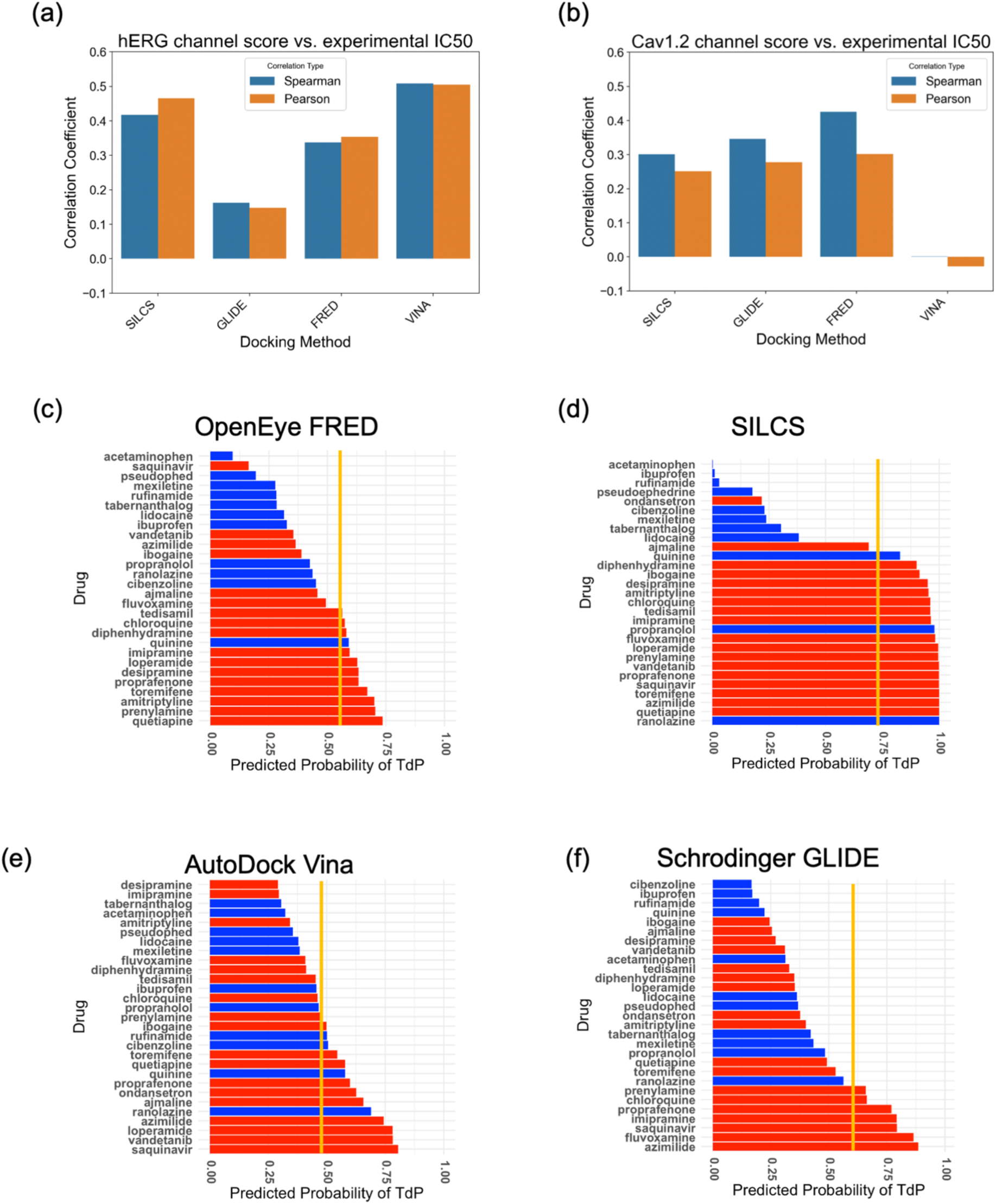
Comparison of docking method scores vs. experimental IC_50_ values. 53 drugs from Kramer et al.^1^ compared to the IC_50_ values from the same study determined from patch-clamp electrophysiology. Both Spearman and Pearson Correlation coefficients reported for both the hERG channel (a) and Ca_V_1.2 (b). (c to e) Logistic Regression models to predict TdP risk based on hERG and Ca_V_1.2 predicted affinities. Docking scores for each method for 53 drugs from Kramer et al.^1^ were used to train the models. Performance of the models was then tested on an independent set of 29 drugs. TdP risk was taken from CredibleMeds where red indicates “*Known risk*” and blue indicates “*Therapeutic Options*.” Optimal threshold shown in yellow.

Attempts to obtain ligand-bound structures of the hERG channel have produced limited results. In 2021, Asai et al. published cryo-EM structures of the hERG channel with and without bound astemizole^27^. A more recent study from the same group provided three additional hERG structures, bound to astemizole, E-4031, and pimozide, three potent hERG-blocking compounds which bind in a very similar site below the selectivity filter.^58^ Given the reasonable correlation of SILCS with both hERG and Cav1.2 vs. experimental data (Figure 2A and B) docked orientations of those compounds generated with that method were examined. Interestingly, SILCS was able to better reproduce the binding pose of these three drugs when no K^+^ ions were placed in the selectivity filter of the channel (Figure S2), with an RMSD of ∼8 Å with ions to ∼4 Å without K^+^. The K^+^ ions are suggested to prevent the positive methylammonium nitrogen (MAMN) fragments from accessing the deep binding site near SF residue Ser624 leading to an absence of favorable positive FragMaps. Given the presence of basic nitrogen atoms on astemizole and pimozide this indicates the importance of the positive FragMaps in accurate modeling of the basic compounds. Accordingly, further model development with the SILCS method used the FragMaps generated in the absence of explicitly K^+^ ions.

Beyond direct regression analysis of the docking scores with respect to the experimental IC_50_ values, we sought to train a simple TdP Risk Classifier using a logistic regression model, as was done previously with experimental IC_50_ values^1^. Model development was performed individually for the 4 docking methods using just the docking scores to hERG and Ca_V_1.2. The logistic models were trained on the 53-drug training set and then tested on an independent set of 29 drugs. The results are shown in Table 1 and panels C to F of Figure 2.

**Table 1:**
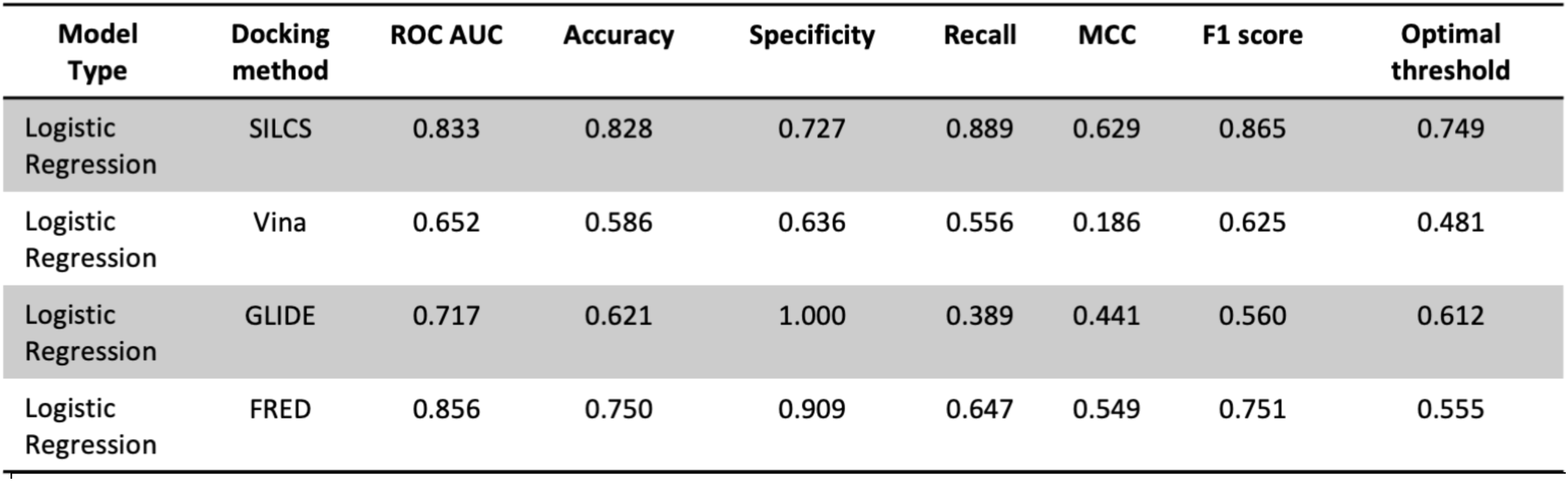
Logistic regression models using different docking programs trained with hERG and Ca_V_1.2 affinity scores.

The model trained on SILCS computed binding affinities resulted in more accurate predictions (Table 1 and Figure 2). The main consideration in the utility of the model was the ability to identify potentially proarrhythmic drugs. Accordingly, we focused on SILCS having the highest Recall, missing only ondansetron as a potentially dangerous drug. However, ondansetron is a frequently prescribed anti-nausea medication that has only moderate risk of TdP^76, 77^ despite CredibleMeds designating it “Known Risk”. Based on these results, further model development focused on the SILCS docking LGFE scores along with selected atomic GFE contributions to those scores.

To potentially further improve model development, we expanded the predicted information available to the classifier model through the inclusion of five additional models of the hERG channel, two additional Ca_V_1.2 channel models, and two models of the Na_V_1.5 channel (see Methods). SILCS docking was performed against all of the models. The docking performance of each model varied considerably (Figures S3-S5). Among the hERG channel models, the best performing model was the low K^+^ structure (Pearson r = 0.74) while the lowest correlation came from the WT structure form Asai et al. (Pearson r = 0.39). With Cav1.2 and Na_V_1.5 poorer correlations versus hERG were obtained though some correlation is evident in all cases.

Further model development to predict TdP Risk incorporated SILCS-based features from the additional structural models. Models were assessed using logistic regression performance on combinations of two or three features as described below. The models were assessed on the original training set using a leave-one-out cross-validation analysis (Figure 3). In addition, a TdP risk model based on the experimental IC values for the three hERG, Cav1.2 and Na_V_1.5 channels was evaluated (H_IC50_, C_IC50_ and N_IC50_, respectively).

**Figure 3:**
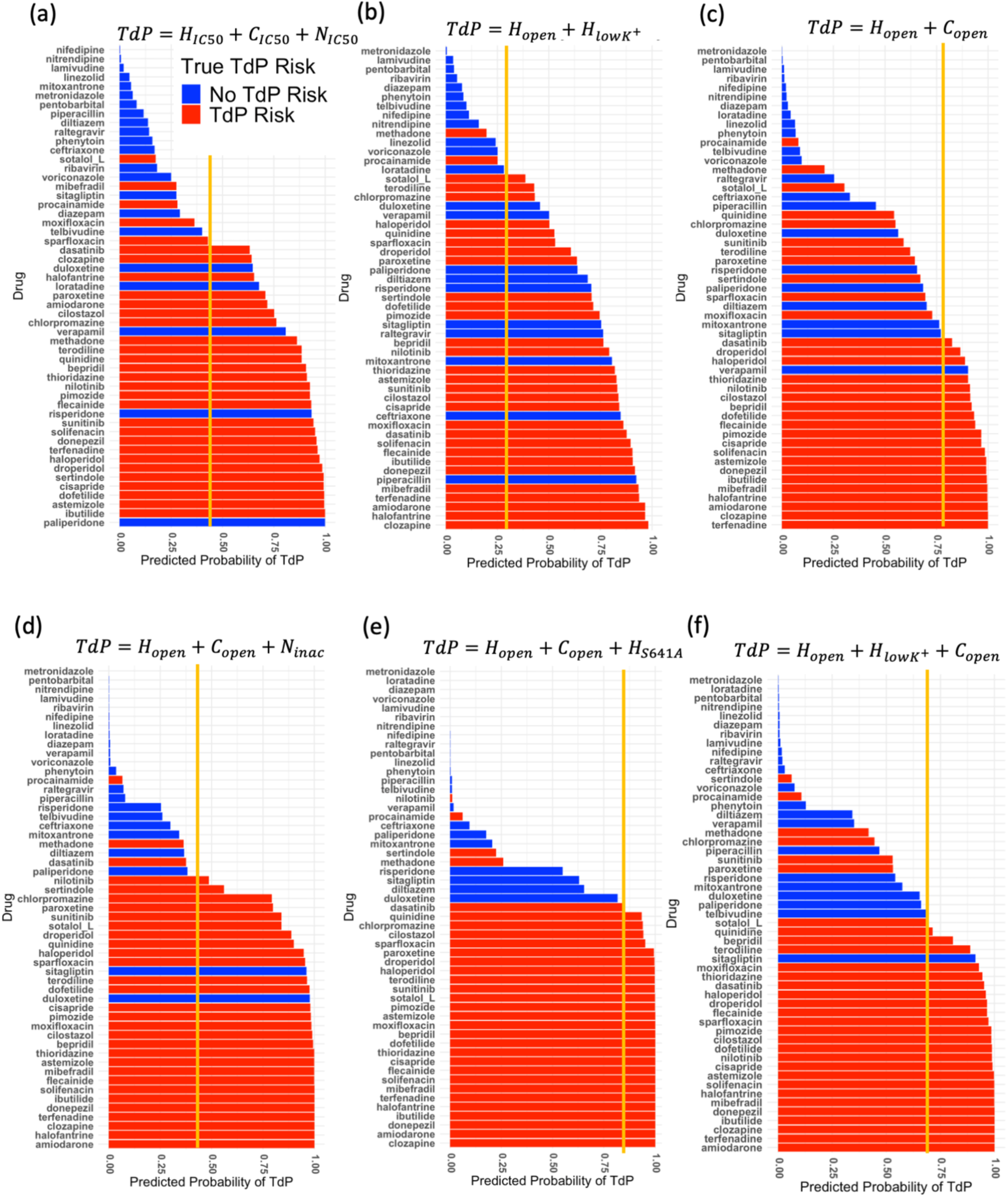
Logistic Regression models trained on SILCS data from additional structures. (a) Model trained with experimental IC_50_ data from Kramer et al.^1^(b) Model with hERG cryo-EM structures with PDB IDs 5VA2 (open) and 9CHQ (low K^+^), (c) hERG 5VA2 and Ca_V_1.2 structure with PDB ID 6AGF, (d) hERG 5VA2, Ca_V_1.2 6AGF, and Na_V_1.5 6LQA structures (e) hERG 5VA2 and Ca_V_1.2 6AGF structure, and hERG S641A mutant models (f) hERG 5VA2, 9CHQ and Ca_V_1.2 6AGF structures. Optimal cutoffs for the risk prediction are shown as yellow lines.

Logistic regression models to generate a predicted probability of TdP for each drug were generated using the SILCS LGFE affinity values to the open (*H_LGFE,open_*) and the low K^+^ “inactivated” states (*H_LGFE,LK_*) of the hERG channel. Generally, the open vs. inactivated hERG model can successfully classify many drugs, with an overall accuracy of ∼77% for the 53-drug test set. Considering each of the two model parameters, *H_open_* and *H_LK_*, only *H_open_* makes a significant contribution to the predictive power of the model (p-value = 0.011). Essentially, adding the additional information of the affinity to the “inactivated” state does not improve the performance of the TdP classifier.

Additional model development incorporated SILCS predicted drug interactions with Ca_V_1.2 channel based on two available cryo-EM structures of hCa_V_1.2 in inactivated conformations (PDBs 8FD7 and 8FHS) as well as a homology model of the open-inactivated state of Ca_V_1.2 channel (that is, Ca_V_1.2 with a more open pore). We observed better model performance for the open-pore model LGFE values, *C_open_* (Figure 3c). Surprisingly, the coefficients for hERG and Ca_V_1.2 are both negative and very similar in magnitude. This hERG vs Ca_V_1.2 model misclassifies ten out of the 53 drugs, with an 81% accuracy (Figure 3c).

Adding as a third term the SILCS LGFE scores of each drug to the inactivated state of Na_V_1.5 led to improved model performance (Figure 3d, Figure S13). Each of the terms in the logistic equation had a significant p-value ( < 0.01) and the overall model accuracy increased to 94%, with just two false positives (duloxetine and sitagliptin) and one false negative (procainamide). Interestingly, this represents an improvement over the accuracy of 83% using the three experimental channel IC_50_ values directly.

Further model development considered the mutant hERG channel models. The S631A hERG mutant experimental structure (PDB 5VA3) did not demonstrate predictive power in model building beyond that achieved by the WT SILCS data (not shown). However, the S641A hERG mutant model, built by introducing the point mutation in silico and performing MD simulation based structural relaxation (see Methods), was useful in combination with the open-state WT hERG channel structure data (Figure 3e). This model achieved a 92% accuracy with only four misclassified drugs: nilotinib, procainamide, sertindole and methadone are all false negatives. The regression coefficients for WT and S641A are both significant and opposite in sign (Figure S11e). This suggests relative drug affinity between the open state WT and S641A mutant hERG structures is predictive of TdP risk.

Finally, using a combination of the SILCS LGFE scores from the hERG open state, the hERG low K^+^ (potentially inactivated) state and the Cav1.2 open state (Figure 3f) yields a strong model. It has a ROC AUC of 0.925 and an 87% overall accuracy. Notably, each of models trained with three sets of docking scores (Figure 3d-f) performed better than the model trained directly on the experimental IC_50_ data which has a ROC AUC of 0.870 and an 83% classification accuracy (Figure 3a).

Based on the results with the smaller data set, we expanded the modeling to include all the drugs classified by CredibleMeds as “Known Risk (TdP+)” for TdP as well as all of the drugs listed on the CredibleMeds page as “Therapeutic Options,” i.e., without TdP risk (TdP-). This resulted in a set of 300 drugs, 80% TdP+ and 20% TdP-. Furthermore, we included 15 physicochemical properties as features in the models, similar to a previous study of hERG block using SILCS.^49^

In addition to a logistic regression classifier, a generalized linear model, we explored three non-linear ML models to improve classification performance: random forest, gradient boosted trees, and a neural network. Each of these models was trained on the larger data set then tested on a reserved set, in a random 80%-20% train-test split. The hyperparameters used to construct these models are listed in Figure S14.

The random forest model had the highest performance, demonstrating the best values for all the metrics evaluated (Table 2), such as a 94% overall model accuracy. The model predicts two false negative results: ondansetron, discussed above, and pentamidine. Interestingly, pentamidine is a drug also misclassified by Kramer et al., and, by them, rationalized as a predictable error since pentamidine is known to disrupt hERG trafficking rather than directly block the pore of the hERG channel.^78, 79^

**Table 2:** TdP Classifier performance summary. Model performance was based.

| Model Type | ROC AUC | Accuracy | Specificity | Recall | Most important positive feature | Most important negative feature | MCC | F1 score |
| --- | --- | --- | --- | --- | --- | --- | --- | --- |
| Logistic Regression | 0.861 | 0.827 | 0.936 | 0.686 | Ca<sub>v</sub>1.2 ACEC | hERG 5VA2 LGFE | 0.492 | 0.591 |
| Random Forest | 0.941 | 0.918 | 0.938 | 0.846 | Ca<sub>v</sub>1.2 GEND | hERG 5VA2 LGFE | 0.763 | 0.815 |
| XGBoost | 0.815 | 0.885 | 0.94 | 0.815 | S641A GENA | hERG 5VA2 LGFE | 0.649 | 0.720 |
| Neural network | 0.760 | 0.803 | 0.896 | 0.462 | Na<sub>v</sub>1.5 MAMC | Na<sub>v</sub>1.5 MAMN | 0.361 | 0.513 |

### 3.3. Ligand design insights directed by ML models

SHAP analysis was leveraged to highlight which features played the strongest role in TdP risk classification. Such information yields insights into the contributions of different features of the drugs to TdP risk, information that can be used to facilitate ligand design (Figure 4a). Notably, SILCS free energy information associated the LGFEs and individual types of functional groups dominated the classifications. The total LGFE binding score of a drug to the hERG channel based on PDB ID 5VA2 (hERG5_LGFE) was the most useful feature at stratifying TdP+ from TdP-drugs. High values of the hERG5_LGFE score, indicating stronger binding to the hERG channel, are consistently observed in classified drugs with a greater TdP risk, as indicated by the top line of Figure 4a. On the other hand, high values of the Nav1.5_MEOO representing drug hydroxyl-group interactions with the sodium channel are predictive of safer drugs.

**Figure 4:**
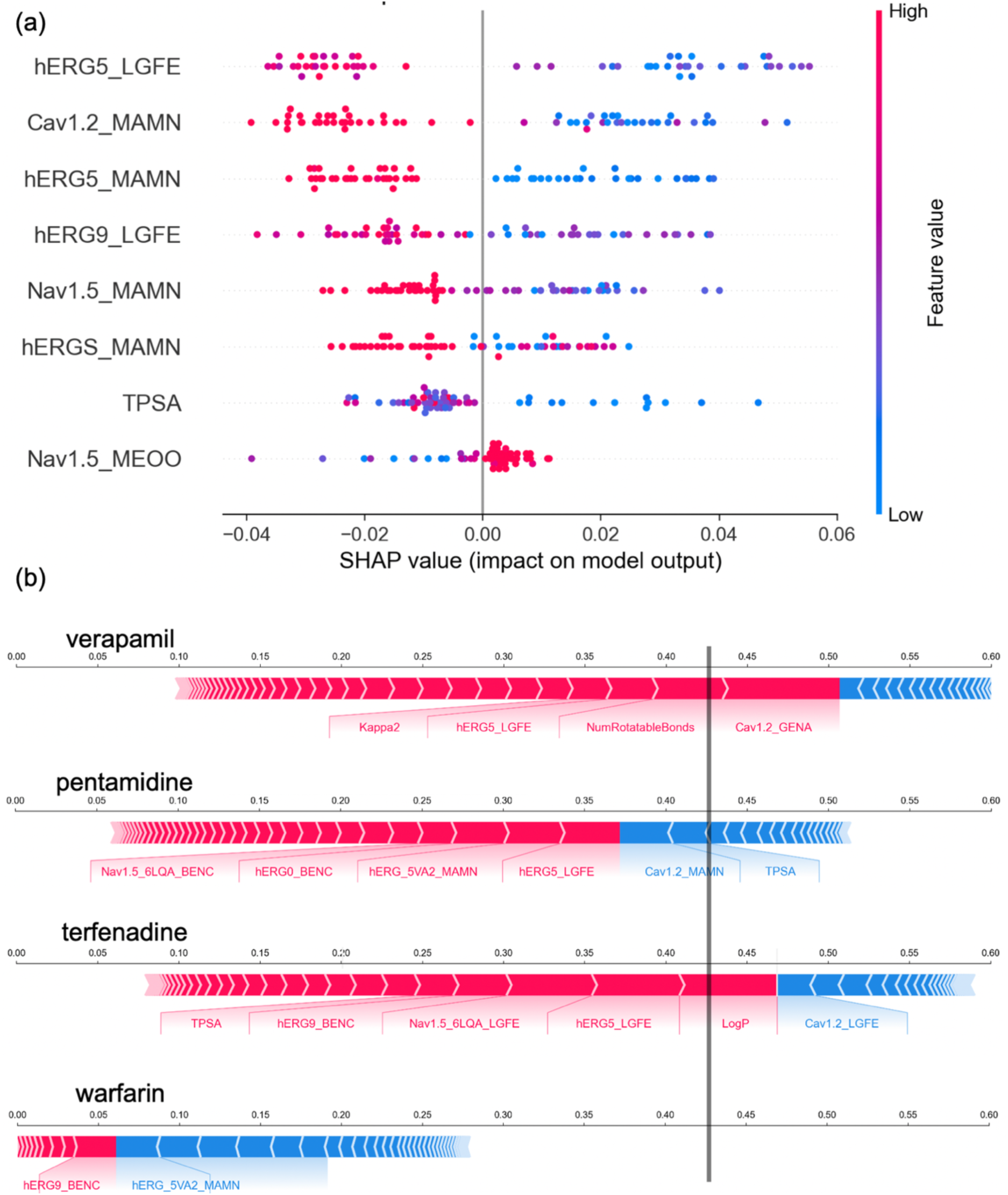
Feature analysis of Random Forest Classifier. (a) SHAP values for eight top features of the model. (b) Force plots for individual drugs indicating how particular model features influenced the predicted probability. The force plots present a balance of features that are predictive of TdP+ as red bars pointing to the right while features the models associate with TdP-drugs are blue bars pointing to the left. The interface of these red and blue features is the predicted probability for a given drug. The prediction threshold is 0.42, shown as the vertical line.

Four drugs provide illustrative examples of the random forest classifier’s performance as highlighted in the force plots of Figure 4b. The force plots present a balance of features that are predictive of TdP+ as red bars pointing to the right while features the models associate with TdP-drugs are blue bars pointing to the left. The interface of these red and blue features is the predicted probability for a given drug. If this interface falls to the right of the prediction threshold of 0.42, the drug is classified as TdP+. Warfarin is a clearly TdP-drug: this correct prediction is driven by low docking scores to the hERG channel. Further, warfarin does not have the cationic amine group that would contribute to the MAMN score. On the other hand, terfenadine is correctly classified as TdP+ due to high hERG binding scores and its LogP score. Here, the model associates highly hydrophobic drugs as more likely to be proarrhythmic. The other two drugs are incorrectly classified: verapamil is associated with low TdP incidence and pentamidine is in fact known to promote TdP. Verapamil is known to be a potent calcium channel blocker which should balance the effects of blocking hERG. The model associates the Ca_V_1.2 generic hydrogen bond acceptor (Ca_V_1.2_GENA) as a proarrhythmic feature. ^80^ Pentamidine receives a TdP-prediction as its hERG binding scores are not high enough to cross the prediction threshold. Based on the model inputs, this is actually the correct prediction since pentamidine likely promotes TdP via disruption of hERG trafficking, a process not addressed by any of the model features.^79^

Notably the LGFE and FragMap information can be directly related to characteristics of a TdP risk. The most direct information comes from specific moieties on a drug being assigned to specific FragMap types. This includes the relationship of hydroxyl groups to MEOO maps and of basic nitrogens to the MAMN maps. In addition, given that that SILCS LGFE scores are based on the summation of individual atomic GFE scores of classified atoms, analysis of the GFE scores summed over different functions groups (GFE values) may be used to identify moieties that make the largest and least contributions to the predicted binding, information that can specifically suggest chemical modifications that can limit TdP risk, which can then be rapidly evaluated using SILCS docking. An example of the information content is shown in Figure 5. Terfenadine risk is largely driven by the basic nitrogen interacting with the hERG channel; replacing this functional group would likely limit TdP risk. Raltegravir, on the other hand, is TdP-(Figure 5a). Hydroxyl groups interacting with the Na_V_1.5 channel are associated with safer drugs (bottom of Figure 4a showing high Nav1.5_MEOO score associated with positive SHAP values). Raltegravir makes more contact with the MEOO map than terfenadine (Figure 5b).

**Figure 5:**
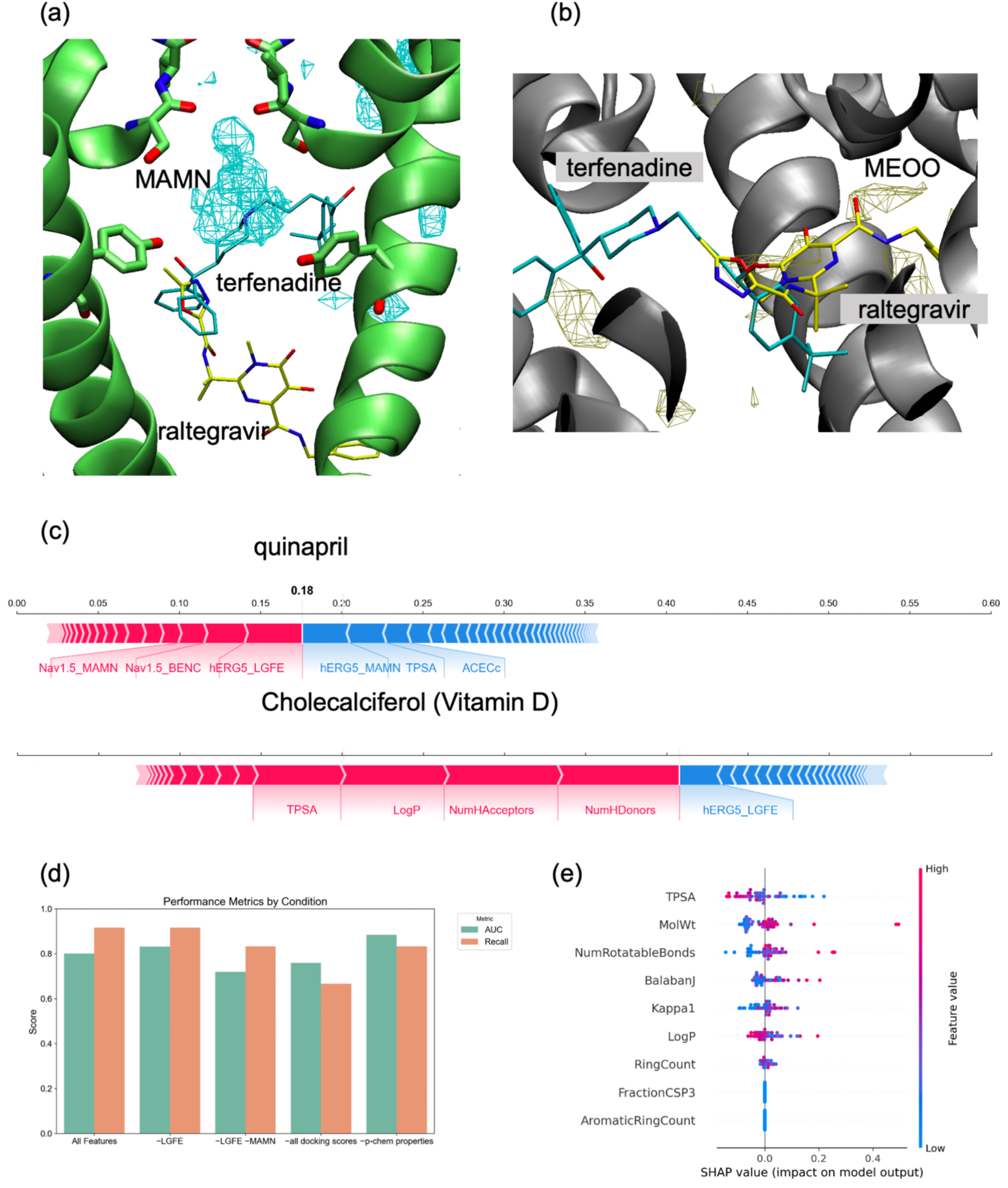
Analysis of important features from Random Forest Classifier. (a) Interactions of the basic Nitrogen with the MAMN FragMap helps discriminate terfenadine from raltegravir, despite the lower overall hERG LGFE of the former. (b) Inclusion of hydroxyl groups that interact with the MEOO FragMaps in Na_V_1.5 is associated with safer drugs. (c) Force plot for quinapril and vitamin D, highlighting contribution of model features to the specific compounds. Predicted probability is shown as the interface between red and blue features. (d) Performance of the model after removing selected features (e.g., -LGFE indicates removing LGFE scores from the model). (e) SHAP analysis highlighting importance of physical properties to RF model.

Looking at the important features from the models, the methylammonium nitrogen (MAMN) score from multiple channel models stood out as a prominent predictor indicate the importance of the presence of a basic nitrogen in ligands. Removing all the MAMN terms leads to a significant drop in ROC AUC and recall score (Figure 5d). One example is highlighted in figure 5a: terfenadine binds much deeper in the hERG channel pore, driven by the overlap of the basic N with the MAMN FragMaps, while raltegravir binds lower in the pore. Raltegravir is correctly classified as a TdP-drug, despite the overall stronger hERG LGFE score.

Notably, removing all the SILCS docking scores also leads to a significant drop in performance, but still leaves a modestly effective classifier. Of the physical properties, only six are useful in the classification (leading to non-zero SHAP values), with topological polar surface area (TPSA) being the most prominent (Figure 5e). Molecular weight and LogP are significant contributors while features like the number of aromatic rings were not meaningfully predictive.

Quinapril and cholecalciferol (vitamin D) provide two illustrative examples of the role of the physical properties of drugs. Quinapril is a clearly TdP-drug that is correctly classified by the model. Despite this, quinapril has a favorable hERG LGFE score. The inclusion of the TPSA parameter in this instance helps keep the predicted risk low (Figure 5c). Cholecalciferol is similarly not known the promote TdP, but the random forest model gives it a predicted probability of 0.41, very close to the threshold. All the top risk factors for this drug are physicochemical properties, acting in opposition to the SILCS docking scores. However, despite these individual successes, the inclusion of the physical properties tends to lead to the classification of multiple safe drugs as TdP+. Further, removing all the physical properties of the model does not significantly affect the model performance (Figure 5d).

## 4. DISCUSSION

Here, we show that structural modeling and docking based on the SILCS-MC method can predict the TdP arrhythmia risk of small molecule drugs in an accurate and high-throughput manner. The approach takes advantage of the precomputed ensemble approach in the SILCS methodology that produces the GFE FragMaps that account for protein flexibility, desolvation, as well as functional group-protein interactions. This allows for rapid docking yielding high accuracy binding affinity predictions in the form of SILCS LGFE scores. This information, which can be partitioned into the contributions of different functional group types, along with simple physical-chemical properties of compounds (e.g., molecular masses, hydrophobicity etc.) can be used as features for development of ML models. The present study yields a Random Forest ML Model that accurately predicts the TdP arrhythmia risk of a range of molecules. Notably, as the approach is based on physics-based energetic information and physical-chemical properties rather than details of the chemical structures themselves, as has been applied previously using the SILCS approach, it is anticipated that the models will be transferrable to a range of chemical space well beyond that used in the training set. Accordingly, it is anticipated that this method will serve as an efficient and significant aid in drug development, representing a significant improvement over current in silico docking methods. The SILCS FragMaps for the studied systems along with the ML models are available at GitHub at https://github.com/kylerouen/tdp_classifiers.

The initial portion of this study involved 4 docking methods (SILCS-MC, Schrodinger Glide, OpenEye FRED, and AutoDock Vina) targeting hERG and Cav1.2 channels. Of the tested methods only the SILCS approach yielded acceptable results in both systems in terms of agreement with experimental IC_50_ values and clinical TdP arrhythmia risks (see Figure 2). Beyond this, we assessed potential state-preferences of drugs by considering SILCS LGFE scores from multiple conformations of the hERG and Ca_V_1.2 channels. Combining the data from several of these structural models leads to a better classifier than considering a single model of each channel (see Figure 3).

Many hERG blocking drugs are positively charged at physiological pH. This leads to the significance of the methylammonium nitrogen (MAMN) GFE binding energy, with cationic drugs attracted to the bottom of the selectivity filter region of the hERG channel, much like K^+^ ions. This is shown directly for terfenadine and raltegravir in Figure 5. Accordingly, it seems that including K^+^ ions in the selectivity filter of the hERG channel can disrupt this high-affinity binding site. Thus, when K^+^ ions are not included in the SILCS simulations, the docking predictions are more accurate. This suggests potent hERG blockers may need to displace K^+^ ions from the bottom of the selectivity filter to reach their preferred binding site.

The relatively low performance of the classifier trained with an open-state hERG model and an inactivated-state hERG model would suggest that any state-dependent hERG channel-drug binding as a predictor of the TdP arrhythmia risk likely takes a back seat to hERG vs. Ca_V_1.2 channel binding as was predicted in some previous studies^1^. The best performing models all included the hERG channel based on PDB 5VA2 with no K^+^ ions in the selectivity filter. Beyond that, we found that several of the other predictor variables are potentially interchangeable. One such set is the open Ca_V_1.2 model and the water-octanol partition coefficient (logP). The logP is relatively anti-correlated with the Ca_V_1.2_open_ score. Accordingly, if you swap Ca_V_1.2_open_ for logP the logistic coefficient flips sign and the model performs very similarly. Another is the Na_V_1.5 quinidine-bound model (based on PDB 6LQA) and the hERG low K^+^ model (based on PDB 9CHQ). These docking scores had a 0.843 correlation. (Figure S7).

In addition to ion channel binding, cellular uptake is required as the drug binding sites are located on the intracellular sides of the channel pores. One way to account for this is in the inclusion of physical-chemical properties such as molecular weight, LogP, and TPSA as features in the classifier model; these features are all associated with cell-membrane uptake and diffusion into ion channel pores. These details, not directly captured by SILCS docking scores, are important contributions to the biological potency beyond the binding affinity of a drug to its target. This is consistent with the observation that making a drug candidate more polar tends to decrease the hERG liability^81^.

Most ML models in this space^7^ focus on predicting whether a compound is a potent hERG blocker or not. In the present study we went beyond this to predict the arrhythmia risk. Further, we aimed to provide insights into physical-chemical properties driving drug binding and the TdP arrhythmia risk rather than present a black box classifier. Such insights are essential to facilitate the rational design of compounds to eliminate TdP arrhythmia liability.

There are multiple classifications for TdP arrhythmia risk (e.g., Redfern et al^82^ and CredibleMeds database: crediblemeds.org). Here, we considered only a binary scheme, assigning drugs with any significant or conditional risk to the same TdP+ category. Others have adopted a more complex five-tiered approach^83^, incorporating different classifications for drugs with few reported TdP arrhythmia incidences or TdP observed only under higher-than-recommended doses. Such an approach that considers intermediate or conditional risk will be the subject of future studies.

In conclusion, we present a pipeline were SILCS docking to multiple protein targets is used to predict the arrhythmia risk of small-molecule drugs. The approach can facilitate the development of new drugs in which an initial screen for cardiac safety with this pipeline would help rapidly narrow the list of drug candidates. In addition, the physics-based approach has the potential to improve the transferability of the models to novel chemical scaffolds and offers molecular-detail information that can greatly facilitate the design of compounds to eliminate arrhythmia risk. Further, this type of approach could be leveraged to design drugs that simultaneously bind multiple targets, e.g., binding of several ion channels to treat atrial fibrillation.^84^

## SUPPORTING INFORMATION

The Supporting Information (SI) is available free of charge at and includes 15 original figures (Figs. S1-S15), which cover details of the ion channels structural models used in the study (Figs. S1, S2, S6), comparison between experimental and computed drug affinities as well as docking parameter optimization details (Figs. S3-S5, S8, S10, S15), TdP arrhythmia prediction machine learning model parameters and performance details (Figs. S7, S9, S11-S14).

## NOTES

A.D.M. is co-founder and CSO of SilcsBio LLC. The other authors do not declare competing financial interests.

## DATA AND SOFTWARE AVAILABILITY

Original simulation data and input files for simulation and analysis have been deposited to GitHub and are publicly available at https://github.com/kylerouen/tdp_classifiers.

## ACKNOWLEDGMENTS

We would like to thank Prof. Heike Wulff for helpful discussions and manuscript comments as well as members of Yarov-Yarovoy and Vorobyov labs for providing some initial protein structural models and valuable suggestions. This work was supported by National Heart, Lung, and Blood Institute (NHLBI) grants F31HL174025 (to K.C.R), R01HL174001, R01HL128537, R01HL152681, and U01HL126273 (to V.Y.-Y. and I.V.), National Institutes of Health (NIH) Common Fund Grant OT2OD026580 (to I.V.), American Heart Association (AHA) Postdoctoral Fellowship Grant 24POST1187017 (to Y.H.), AHA Career Development Award grants 25CDA1450133 (to Y.H.) and 19CDA34770101 (to I.V.), UC Davis Chemical Biology Program fellowship supported in part by NIGMS Institutional Training Grant T32GM136597 (to K.C.R.), UC Davis T32 Predoctoral Training in Basic and Translational Cardiovascular Medicine fellowship supported in part by NHLBI Institutional Training Grant T32HL086350 (to K.C.R.), Oracle for Research fellowship (to I.V.), and NIGMS R35 GM131710 grant (to A.D.M.). Computer allocations were provided through Advanced Cyberinfrastructure Coordination Ecosystem: Services & Support (ACCESS) and Extreme Science and Engineering Discovery Environment (XSEDE) grants MCB170095 (to I.V. and V.Y.-Y.), Texas Advanced Computing Center (TACC) Pathways Allocation MCB20010 (to I.V. and V.Y.-Y.), Oracle cloud credits award (to I.V.).

