## Supplemental Information for "Prediction of TdP Arrhythmia Risk Through Molecular Simulations of Conformation-specific Drug Interactions with the hERG K^+^, Na_V_1.5, and Ca_V_1.2 Channels"

### for

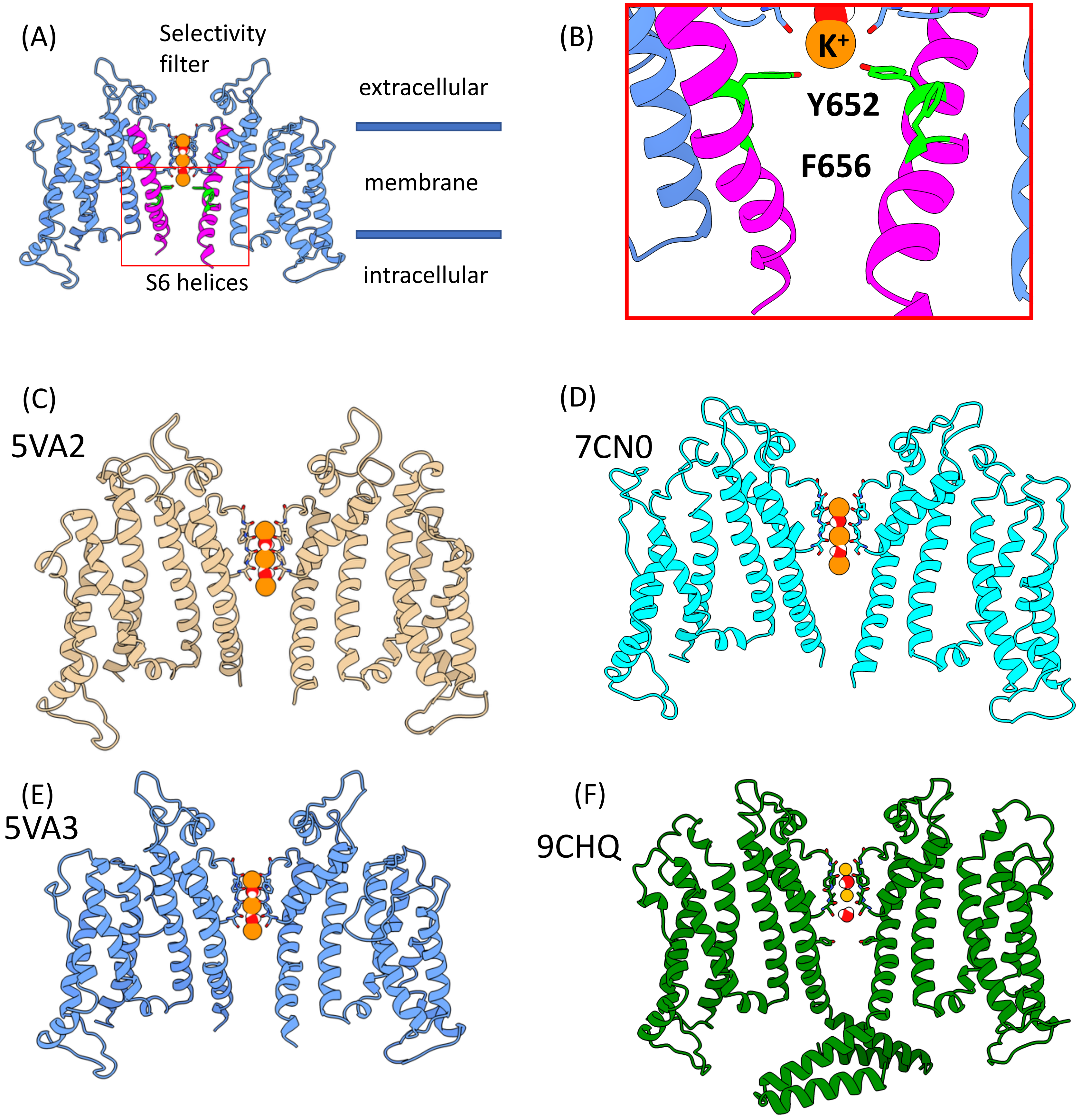

***Figure S1:*** **Structural models of the hERG ($K_V$11.1) channel.** (A) Structure of the channel with two opposite subunits relative to the lipid membrane position. (B) Notable drug binding residues Y652, F656 relative to the $K^+$ ion binding site. (C) Structure model of the open state based on PDB 5VA2 (D) Structural model of the inactivated state based on PDB 7CN0. (E) Structural model of the non-inactivating mutant based on PDB 5VA3. (F) Structural model of the low K+ WT hERG channel based on PDB 9CHQ.

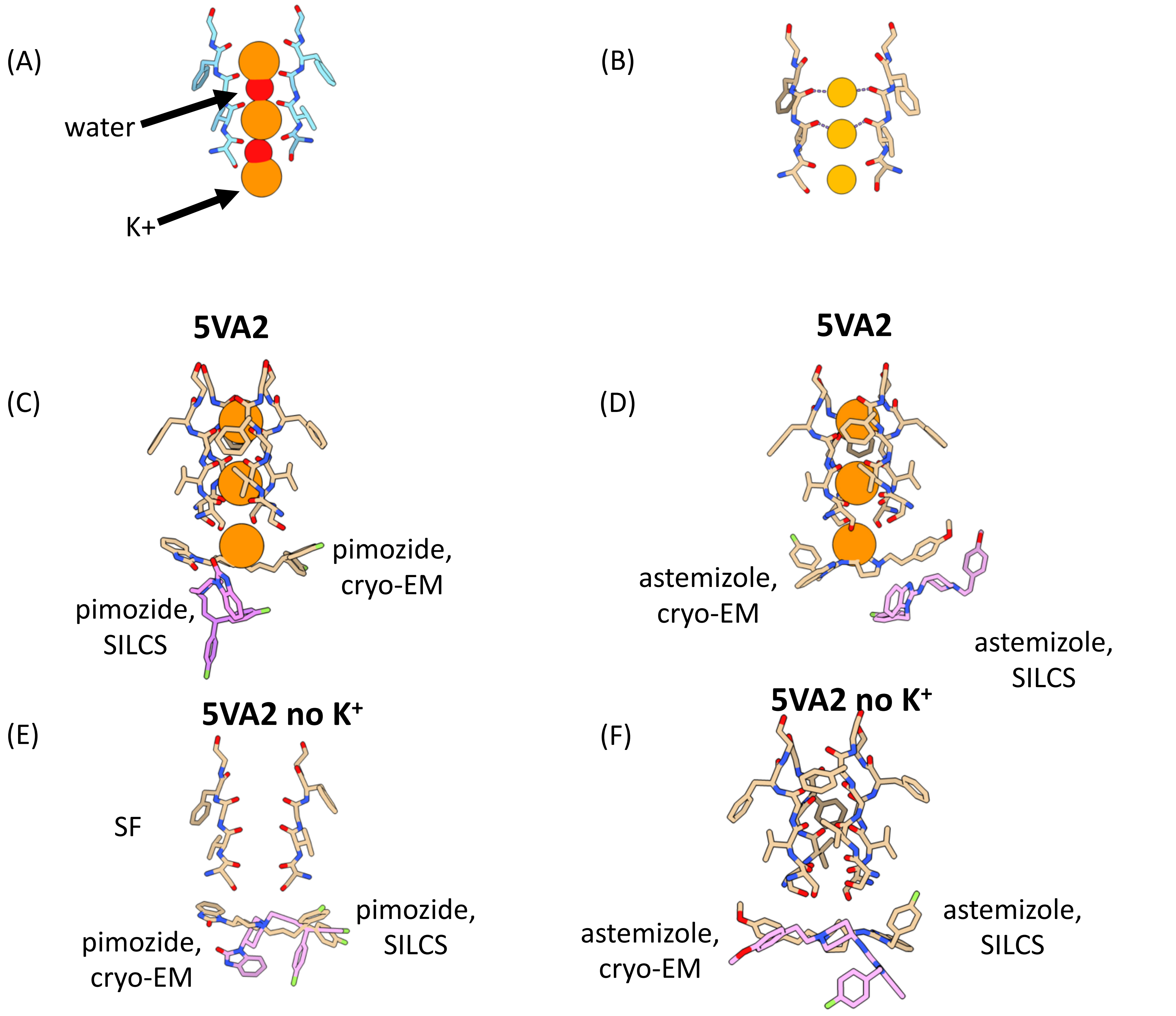

***Figure S2*: hERG channel - drug interactions.** (A) Ion and SF waters used with 5VA2. (B) Ions from PDB 7CN1. (C-F) SILCS docked (pink) and experimental (brown) orientations of (C) pimozide: RMSD (non hydrogens) = 8.8 Å; (D) astemizole: RMSD = 10.6 Å; (E) pimozide: RMSD = 4.6 Å. (F) astemizole: RMSD = 8.3 Å. (C) and (D) show docking poses with $K^+$ ions while (E) and (F) show poses without $K^+$ ions which leads to lower RMSDs with respect to cryo-EM binding poses (8ZYO for astemizole and 8ZYQ for pimozide).

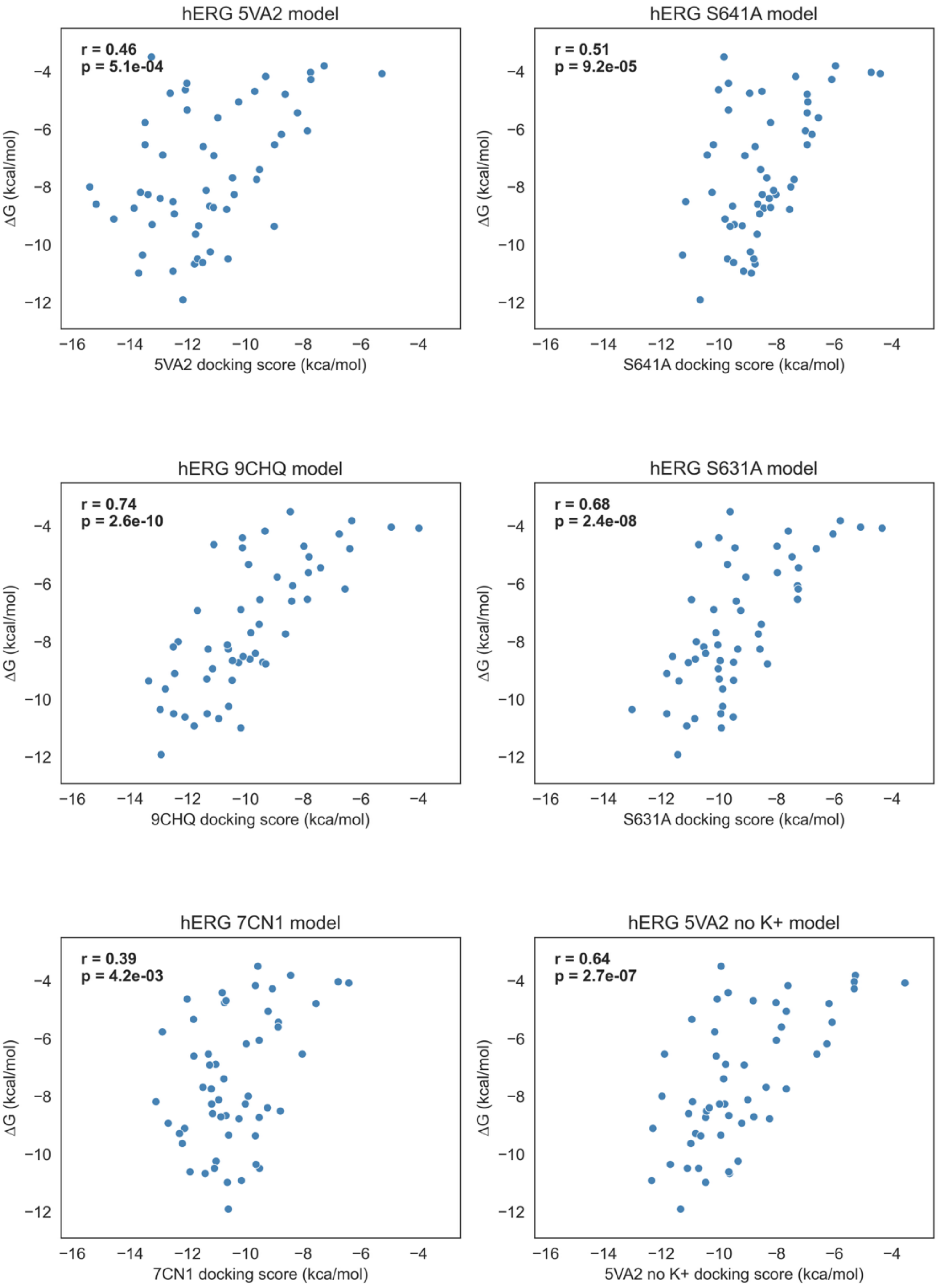

***Figure S3:*** **Correlation of SILCS LGFE scores and experimental data using different hERG channel models**. Experimental DG values from the reported IC50 values are shown on the Y axes and SILCS LGFE docking scores are shown on the X axes.

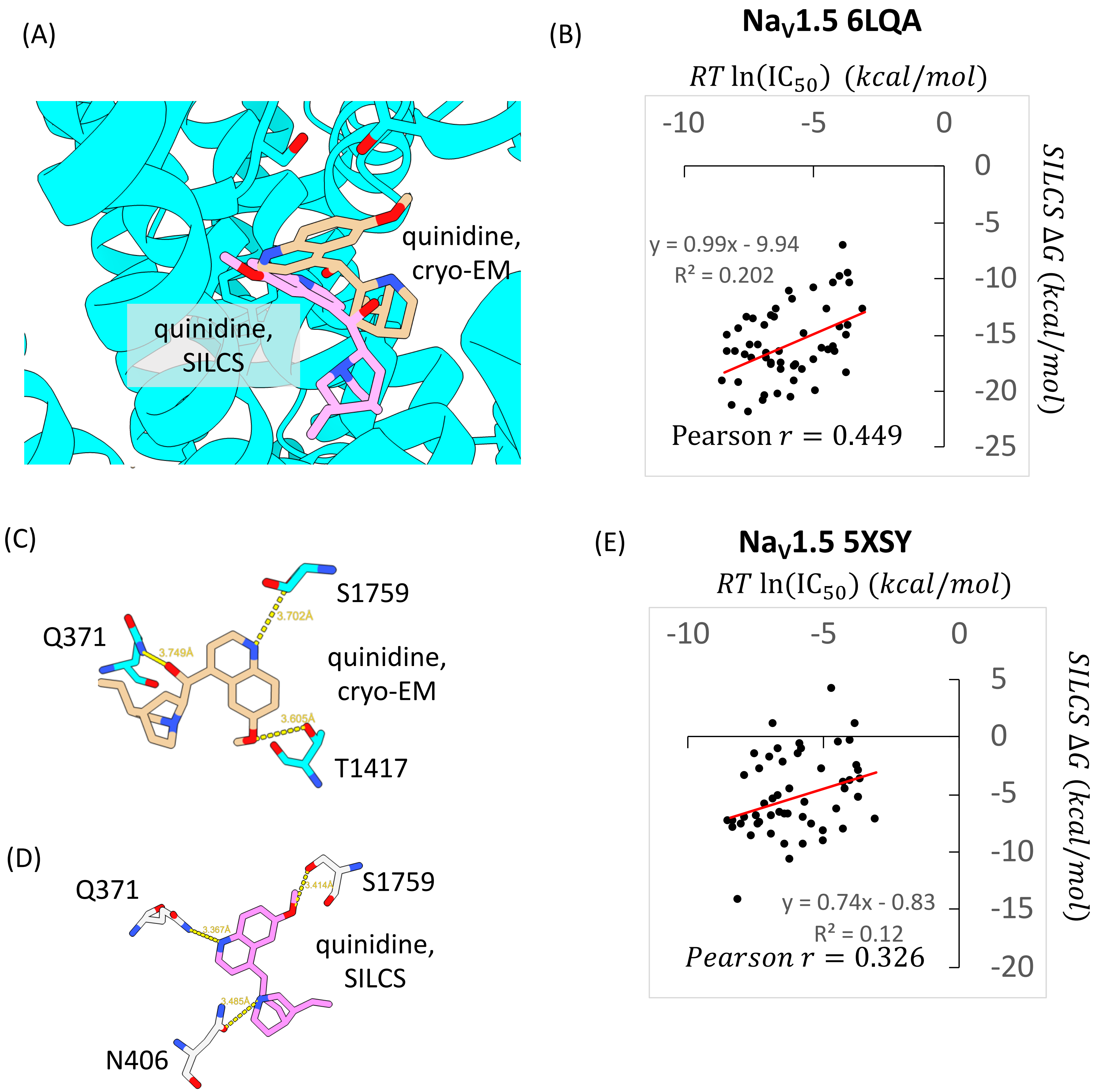

***Figure S4:* Na$_V$1.5 channel - drug interactions.** (A) SILCS quinidine pose (pink) compared to cryo-EM quinidine pose (brown) from 6LQA, RMSD (non-hydrogen) = 5.22 Å. (B) Correlation between experimental $IC_{50}$ from Kramer et al. and SILCS LGFE scores for 53 drug test set for PDB 6LQA. IC50 values converted to kcal/mol as –RTln(IC50). (C) Quinidine contacts from SILCS include a new contact with N406 and the tertiary amine instead of T1417 and the QDN hydroxyl. (D) Polar contacts from cryo-EM show no contact with tertiary amine but hydroxyl hydrogen bonds with T1417. (E) Correlation between experimental $IC_{50}$ from Kramer et al. and SILCS docking score for 53 drug test set for homology model based on PDB 5XSY.

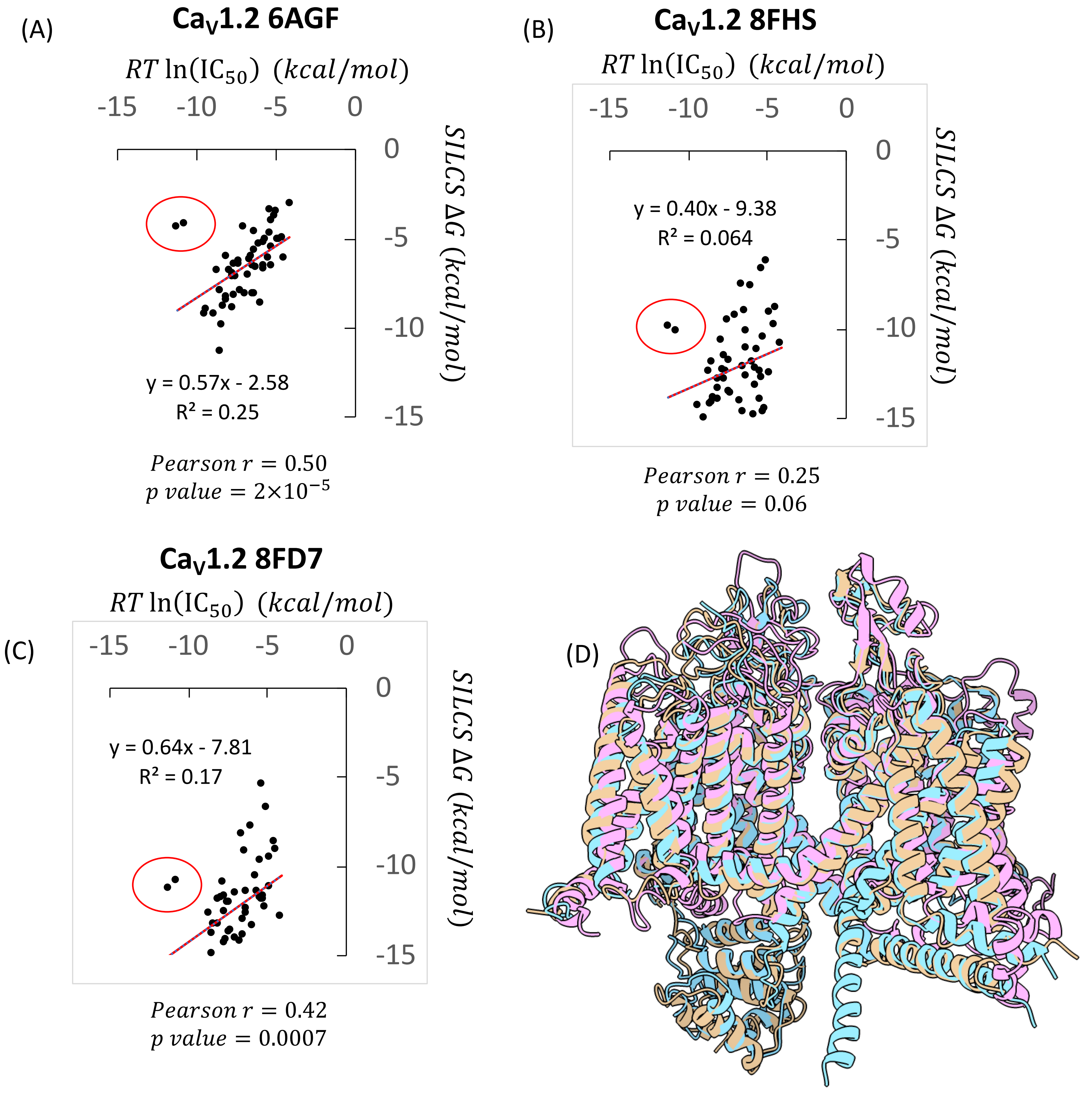

***Figure S5:*** **Docking of 55 hERG-blocking drugs compared to experimental affinity from Kramer et al.** (A) Experimental affinity vs docking to the $Ca_V$1.2 homology model based on PDB 6AGF. (B) Experimental affinity vs docking to the $Ca_V$1.2 8FHS model. (C) Experimental affinity vs docking to the $Ca_V$1.2 8FD7 model. Scores for nifedipine and nitrendipine are circled in each panel. (D) Alignment of PDB 6AGF (pink), 8FHS (blue), and 8FD7 (pink) structures.

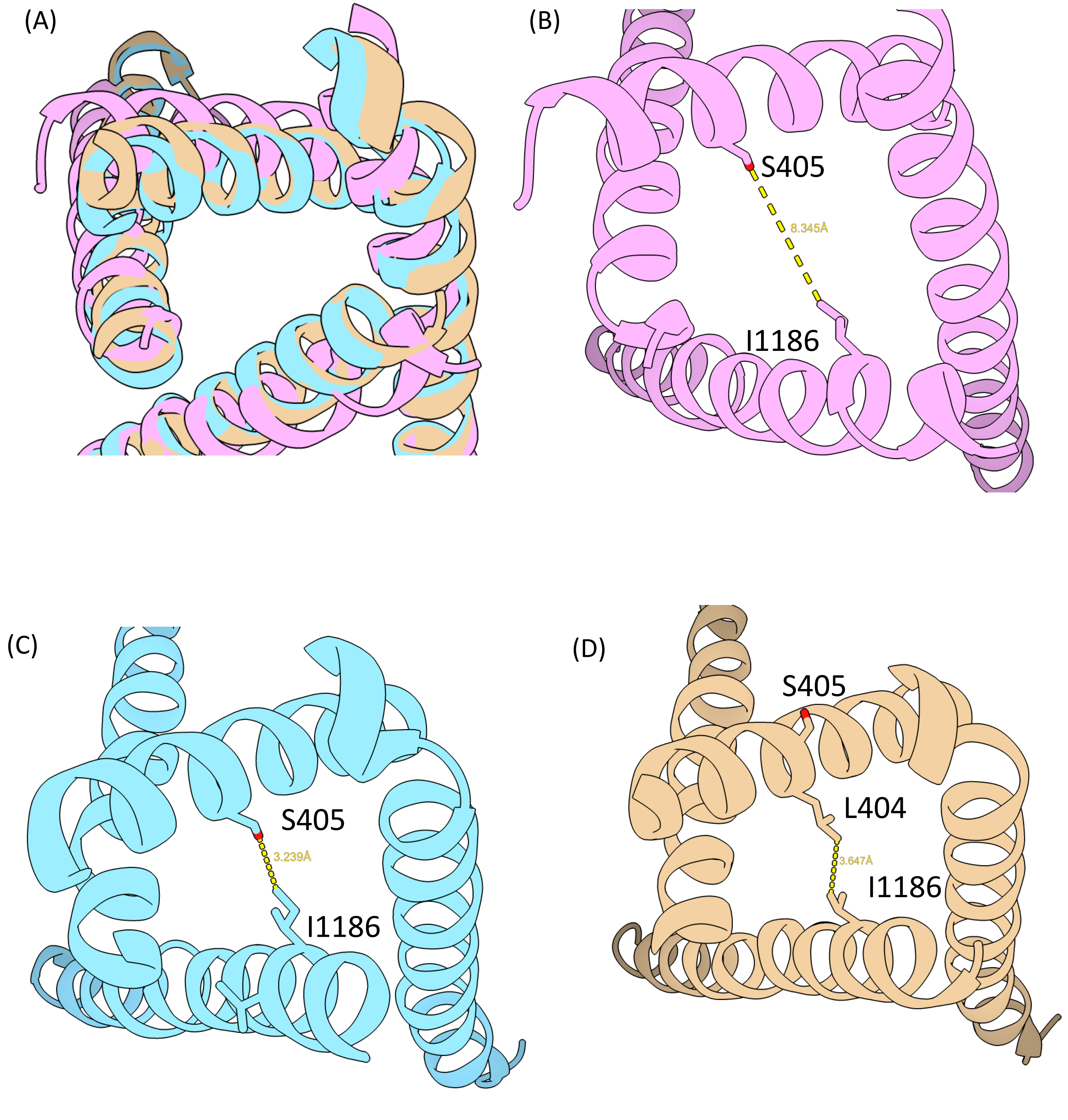

***Figure S6:*** **Cav1.2 channel structural models**. (A) Overlay of S6 helices for 3 Cav1.2 models including (B) Homology model based on PDB 6AGF (pink), (C) 8FHS model (blue) and (D) 8FD7 model (brown).

(A)

$$TdP = H_{open} + C_{open} + H_{S641A}$$

$$TdP = H_{open} + logP + H_{S641A}$$

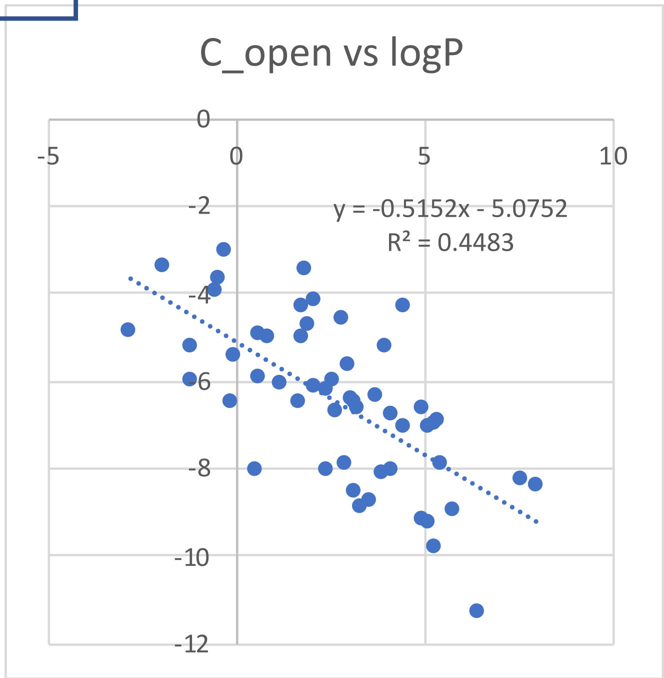

(B)

$$TdP = H_{open} + C_{open} + H_{LK+}$$

$$TdP = H_{open} + C_{open} + N_{inac}$$

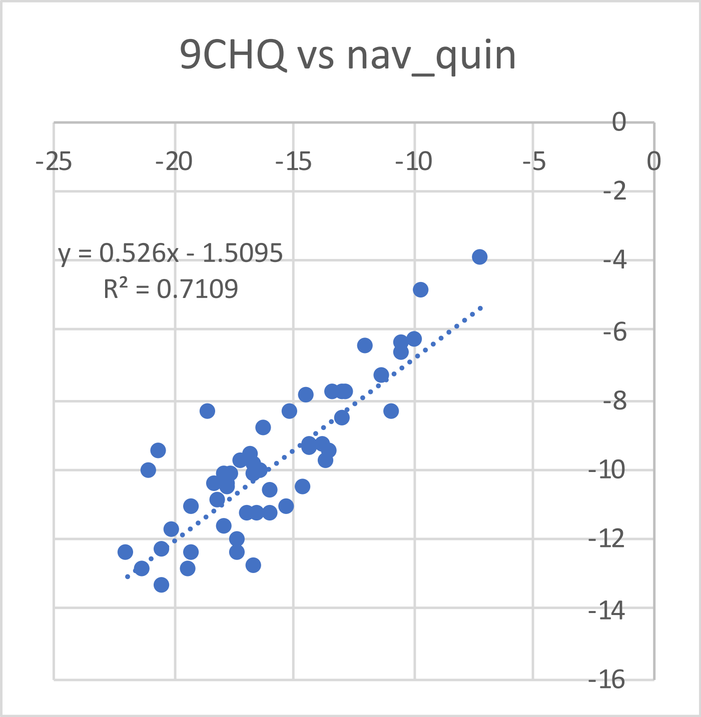

***Figure S7:*** **Correlation between different TdP risk prediction model scores.** (A) Correlation between open-state Cav1.2 LGFE scores ($C_{open}$) and LogP and (B) correlation between between low K+ hERG LGFE scores ($H_{LK}$) and inactivated Nav1.5 LGFE score ($N_{inac}$).

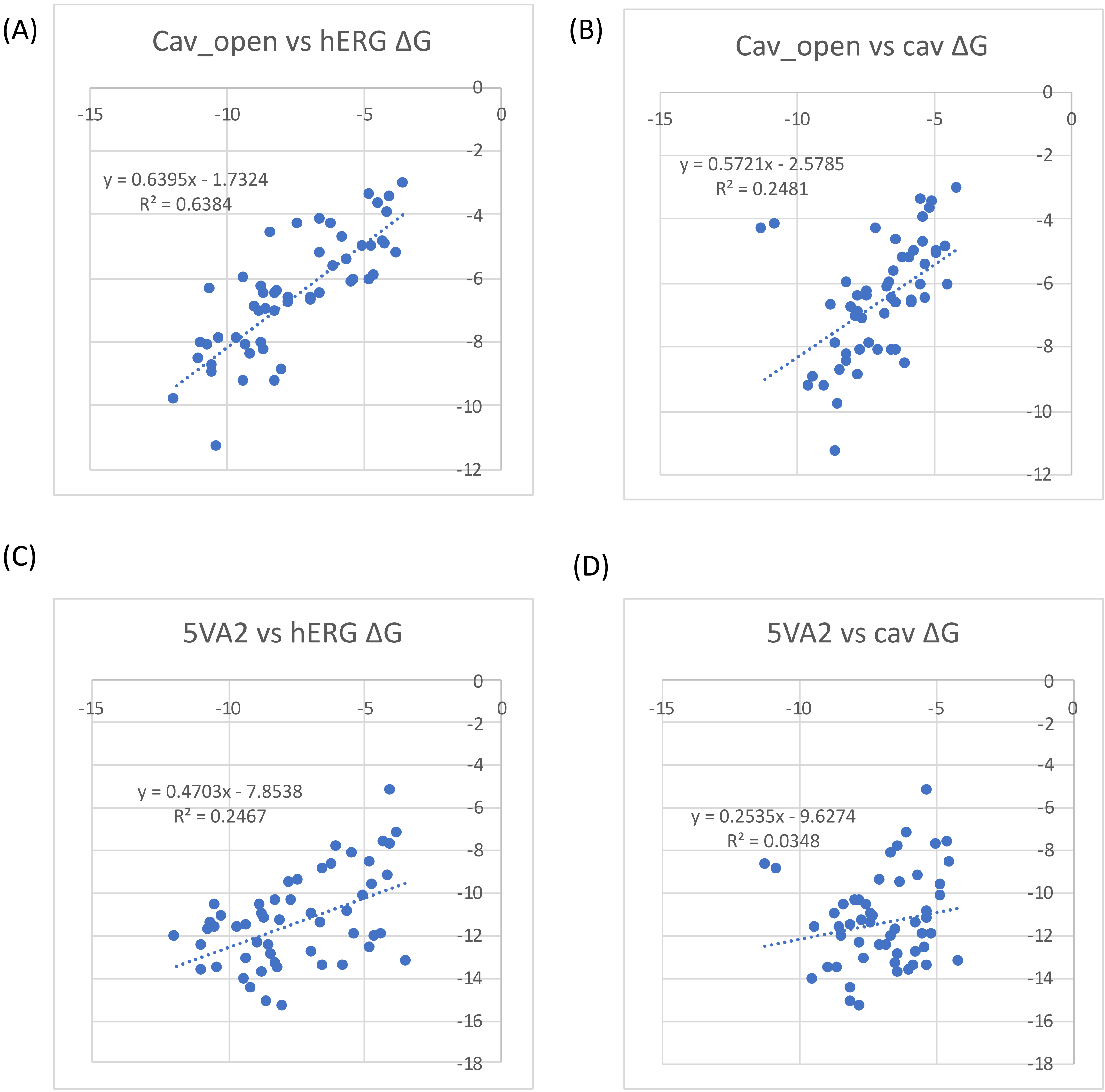

***Figure S8:*** **Correlation between SILCS LGFE scores and experimental ΔG values.** (A) Open Cav1.2 docking vs experimental hERG ΔG (B) Open Cav1.2 LGFE scores vs experimental Cav1.2 ΔG (C) Open hERG LGFE scores vs experimental hERG ΔG and (D) Open hERG LGFE scores vs experimental Cav1.2 ΔG.

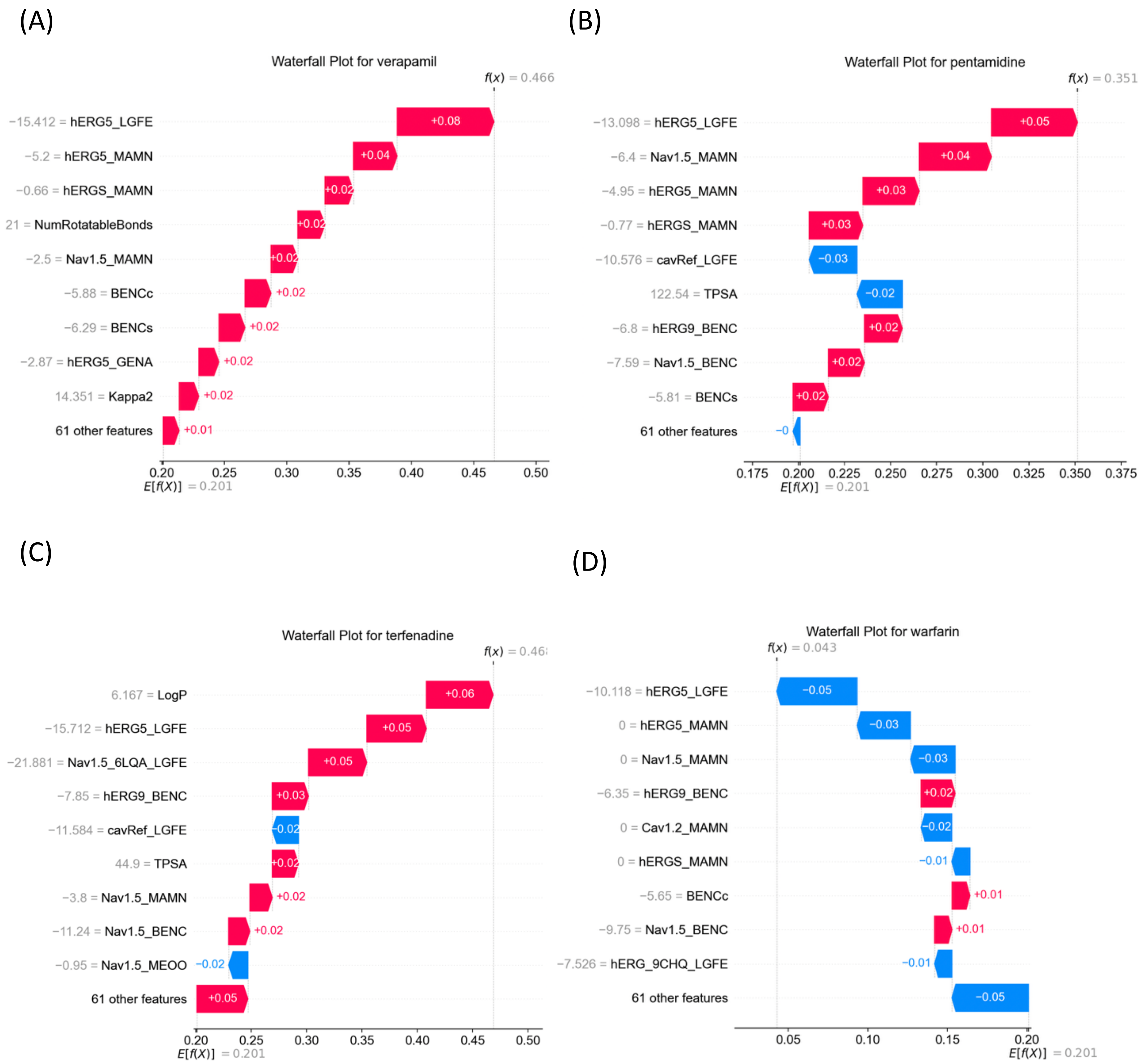

***Figure S9:*** **Waterfall plots showing feature importance from Random Forest classifier** for (a) verapamil (b) pentamidine (c) terfenadine and (d) warfarin.

(a)

| Param Set | Gauss1 | Gauss2 | Repulsion | Hydrophobic | Hydrogen Bond | Rotation |
|---|---|---|---|---|---|---|
| Default | −0.0356 | −0.00516 | 0.840 | -0.0351 | -0.587 | 0.0585 |
| Pham et al. | -0.049811 | -0.007218 | 0.756221 | -0.031562 | -0.469951 | 0.025722 |
| hERG optimized | -0.049811 | -0.007218 | 0.756221 | -0.04559 | -0.469951 | 0.025722 |

(b)

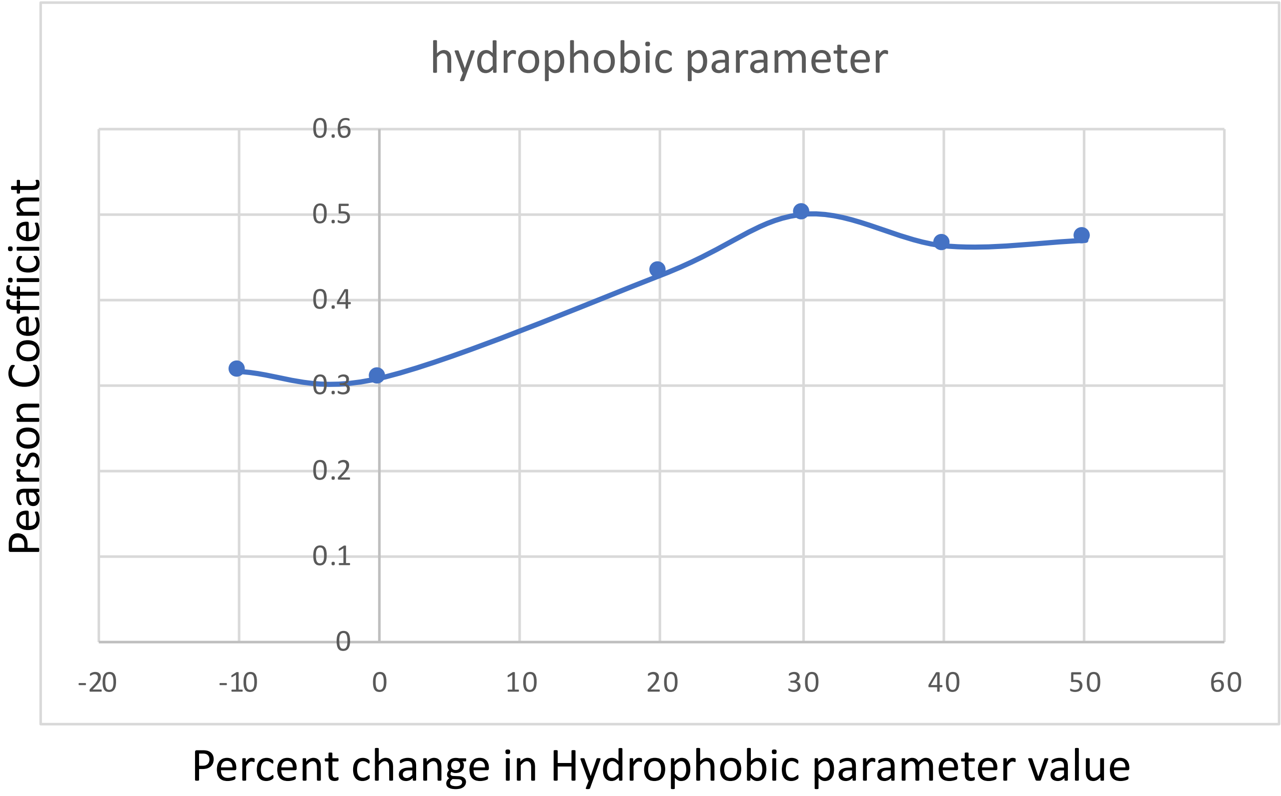

***Figure S10:*** **AutoDock Vina parameter optimization.** (a) optimized parameter set for the hERG channel with increased hydrophobic weight and (b) correlation of test drugs has a function of hydrophobic parameter.

| (a) | Coefficient | p-value |
|---|---|---|
| hERG5 | -0.760 | 0.005 |
| hERG9 | -0.166 | 0.5 |
| intercept | -9.774 | 0.001 |

| (b) | Coefficient | p-value |
|---|---|---|
| hERG5 | -0.982 | 0.006 |
| Cav1.2 open | -0.932 | 0.004 |
| intercept | -16.502 | 0.002 |

| (c) | Coefficient | p-value |
|---|---|---|
| hERG5 | -1.62 | 0.003 |
| hERG9 | -0.804 | 0.03 |
| hERG S641A | 1.78 | 0.01 |
| intercept | -10.243 | 0.005 |

| (d) | Coefficient | p-value |
|---|---|---|
| hERG5 | -3.139 | 0.007 |
| Nav1.5 | 1.164 | 0.01 |
| Cav1.2 open | -2.108 | 0.003 |
| intercept | -29.204 | 0.004 |

| (e) | Coefficient | p-value |
|---|---|---|
| hERG5 | -5.124 | 0.03 |
| hERG S641A | 4.636 | 0.03 |
| Cav1.2 open | -3.468 | 0.04 |
| intercept | -37.776 | 0.05 |

| (f) | Coefficient | p-value |
|---|---|---|
| hERG5 | -1.868 | 0.003 |
| hERG9 | 1.281 | 0.02 |
| Cav1.2 open | -2.228 | 0.005 |
| intercept | -21.650 | 0.004 |

***Figure S11:*** **Terms from Logistic Regression Analysis using leave-one-out cross validation on 53-drug training set.** Model performance shown in main-text Figure 3.

(a)

| Model terms | ROC AUC | Sensitivity | Specificity | Accuracy |
|---|---|---|---|---|
| hERG IC50 | 0.852 | 0.774 | 0.864 | 0.811 |
| hERG LGFE | 0.811 | 0.970 | 0.591 | 0.811 |
| hERG + Cav IC50 | 0.871 | 0.818 | 0.880 | 0.849 |
| hERG + Cav LGFE | 0.886 | 0.645 | 0.955 | 0.774 |
| hERG + Nav + Cav IC50 | 0.870 | 0.871 | 0.773 | 0.830 |
| hERG + Nav + Cav LGFE | 0.943 | 0.903 | 0.909 | 0.906 |

***Figure S12:* Model performance from Logistic Regression Analysis using leave-one-out cross validation on 53-drug training set.** IC50 indicates experimental values from Kramer et al. and LGFE indicates SILCS docking score to the respective ion channels.

| Topological Index | Description |
|---|---|
| Balaban J Index ($J$) | $$J=\frac{N}{M-N+2}\sum_{bonds\ i<j}\left(s_i s_j\right)^{-1/2}$$ $N$=number of heavy atoms, $M$=number of bonds. $s_i$=sum of the distances from atom $i$ to every other atom (measured in number of bonds). High values of $J$ indicate linear molecules while low values indicate more branched or cyclized molecules. |
| Kappa1 ($\kappa_1$) | $$\kappa_1=\frac{(N-1)^2}{M^2}$$ High values indicate long, linear molecules with minimal branching. |
| Kappa2 ($\kappa_2$) | $$\kappa_2=\frac{(N-1)(N-2)^2}{P_2^2}$$ $P_2$=number of paths with length two. High values indicate a molecule a high degree of branching. |
| Kappa3 ($\kappa_3$) | $$\kappa_3=\frac{(N-1)(N-3)^2}{P_3^2}$$ $P_3$=number of paths with length three. High values indicate a molecule with a high number of cycles/rings. |
| Bertz Complexity Index ($C_T$) | $$C_T=\ln\left[\prod_{i=1}^{N}(d_i)!\right]$$ $d_i$=number of atoms directly bonded to atom $i$. High values of $C_T$ indicate more complex molecules with a high degree of branching and/or cycles. |

***Figure S13:*** **Description of topological indices used as model features.**

| ML algorithm | hyperparameters | Value range tested | Final value |
|---|---|---|---|
| Random Forest (RF) | n_estimators | (50, 300) | 74 |
| | max_depth | (3, 30) | 24 |
| | min_samples_split | (2, 20) | 3 |
| | min_samples_leaf | (1, 20) | 1 |
| XGBoost (XGB) | n_estimators | (50, 300) | 193 |
| | max_depth | (3, 10) | 9 |
| | learning_rate | (0.01, 0.2) | 0.09 |
| | subsample | (0.6, 1) | 0.68 |
| | colsample_bytree | (0.6, 1) | 0.77 |
| | reg_alpha | (0.0, 1) | 0.19 |
| | reg_lambda | (1, 3) | 1.05 |
| Neural Network (NN) | hidden1 | (32, 128) | 124 |
| | hidden2 | (16, 64) | 124 |
| | dropout | (0.1, 0.5) | 0.448 |
| | learning_rate | (0.0001, 0.01) | 0.0073 |
| | weight_decay | (0.000001, 0.01) | 0.00085 |
| | n_epochs | 100 | 100 |

***Figure S14:*** **Hyperparameters from ML classifier models.** Values were varied uniformly within the range listed by the Optuna software package to optimize the ROC AUC of each model.

(A) (B) (C)

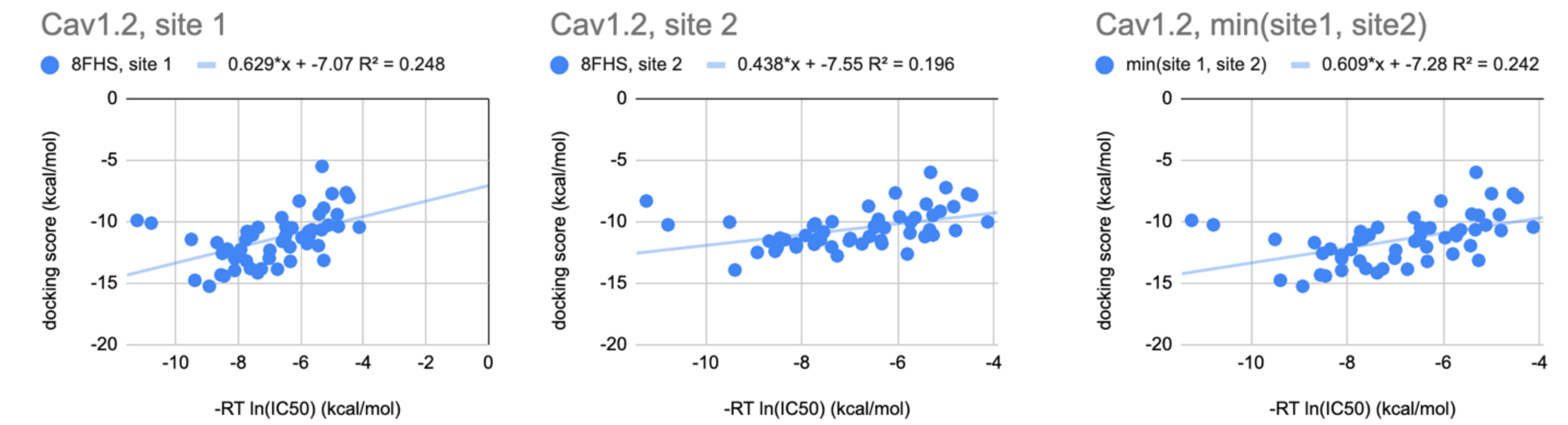

(D)

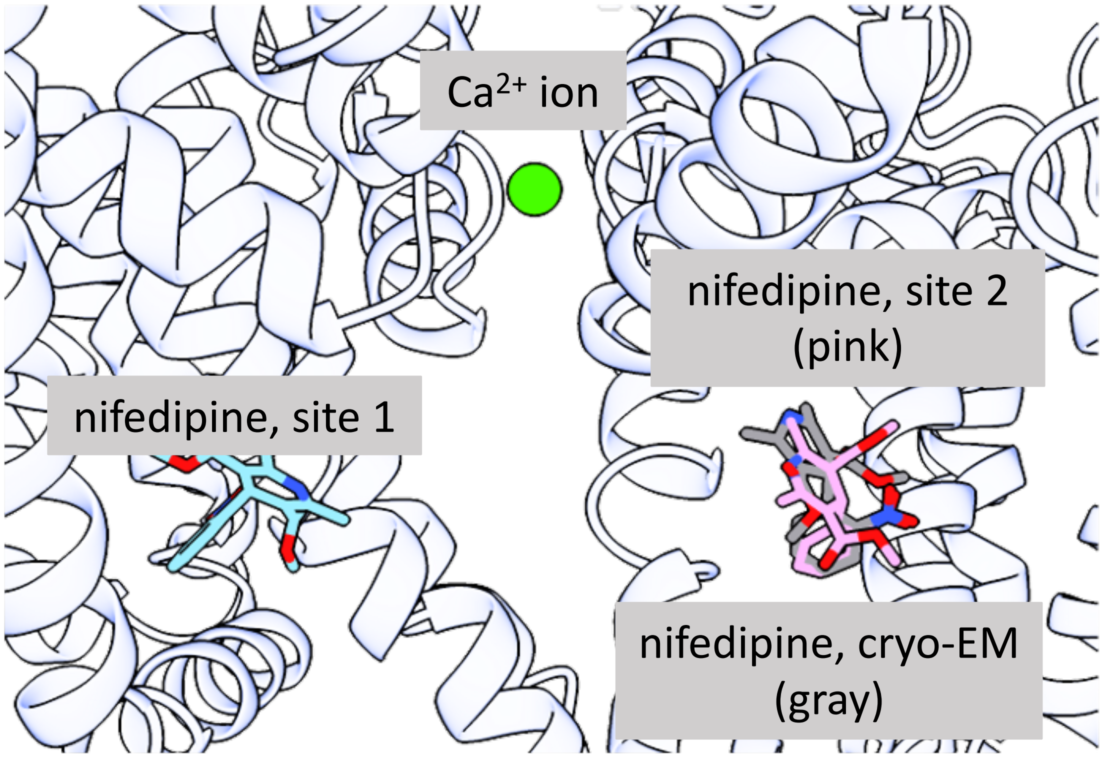

| | docking score (kcal/mol) | |
|---|---|---|
| **drug** | **Site 1** | **Site 2** |
| nifedipine | -9.901 | -8.321 |
| nitrendipine | -10.119 | -10.262 |

***Figure S15:*** **Ca$_V$1.2 channel drug docking results using OpenEye FRED.** (A) OpenEye docking scores for the 53 drugs from Kramer et al. to Site 1 of Ca$_V$1.2 (PDB 8FHS). Site 1 is the result of broad docking within the entire intracellular cavity which results in a minimum energy pose near the fenestration between domains I and II. (B) Site 2 is the result of a narrower search area, focused on the dihydropyridine (DHP) binding site, the fenestration between domains III and IV. (C) Docking score correlation taking the lower value from site 1 and 2 vs. experimental IC$_{50}$. (D) Nifedipine docking poses produced by OpenEye FRED in two different fenestration sites. Cryo-EM pose of nifedipine from the rabbit Ca$_V$1.1 structure (PDB 6JP5). The docking score for nitrendipine, a related DHP antagonist, also does not differ significantly between the two sites.
